# Percolation-inspired criticality in complement activation: universal scaling and transport-limited complement surface amplification

**DOI:** 10.64898/2026.08.14.744667

**Authors:** Stephanie Monson, Sahil Kulkarni, Jacob Meyerson, Jacob Brenner, Ravi Radhakrishnan

## Abstract

The collective spatial phenomenon of complement protein opsonization on nanoparticle surfaces is a key component of the immune response to viruses, engineered nanoparticles, and diseased cells. Recent work showed this opsonization follows a sharp, percolation-like transition versus the spacing d between surface-bound attachment sites, leaving two open questions: 1) whether the transition exhibits hallmarks of true criticality, such as diverging susceptibility, and 2) whether it can be distinguished from an alternative first-order cooperative (Hill-type) process producing an equally sharp threshold without true criticality. Here, we resolve both questions using a hierarchical statistical-mechanics treatment spanning stochastic, mean-field, and spatial reaction-diffusion models. The variance of two order parameters, peak complement activity and activation lifetime, diverges near threshold and sharpens systematically with system size, the defining signature of a critical point rather than a smooth cooperative response. Extending the analysis across site spacing and intrinsic kinetic rate constants traces a two-dimensional locus of critical points with consistent critical exponents throughout, establishing a single, robust universality class. The mean-field dynamic exponent for activation lifetime agrees quantitatively with the exact value predicted for the general epidemic process. Finally, a reaction-diffusion model of the nanoparticle surface shows the critical locus is set by a diffusion-limited length scale, establishing complement percolation as a fundamentally transport-limited surface reaction. These results place complement activation within the percolation universality class and identify the physical parameters, diffusion, catalysis, and decay, that govern its critical threshold, with direct implications for rational design of complement-evading nanomaterials, immunology, and evolutionary biology.

**Significance Statement:** The immune system must decide, within seconds, whether a foreign surface is dangerous, not through complex signaling, but through collective phenomena playing out in space. We show complement activation, the cascade cells use to tag threats for destruction, is not a steep biochemical response but a true critical phenomenon: the mathematical class of transition governing percolation, epidemic spreading, and magnetism. A handful of physical quantities, how fast molecules diffuse, catalyze, and decay, set a universal threshold where activation switches from quiescent to triggered, exploiting a deep organizing principle of physics and revealing immune recognition as predictable from statistical mechanics rather than molecular detail alone.

## Introduction

Spatial collective transitions in biochemical networks are the basis of many biological functions. One example is the spreading of complement proteins on nanoscale surfaces, such as those of engineered nanoparticles, viruses, and extracellular vesicles. The complement system is a part of the innate immune system, which defends against pathogens and other foreign entities. Complement response to engineered nanomaterials results in poor biodistribution and efficacy of therapeutics, rejection of transplants, and, in severe cases, potentially deadly allergic reactions^1–7^. There have been many calls from the nanomaterials community for better understanding the relationship between immune reactions to nanomaterials and nanomaterial properties to design safer and more effective nanomaterials. On the other hand, complement response to viruses and other foreign biological entities, like cancer cells, is desirable; however, viruses and cancer cells have adapted strategies to avoid complement recognition^8,9^. Furthermore, excess complement coating of nanoparticle surfaces is implicated in diseases like Alzheimer’s disease and age-related macular degeneration^10–12^. Characterizing the collective transition that occurs between non-complement-coated and complement-coated surfaces would be of great benefit for understanding health and disease.

The relationship between complement and nanoparticle surfaces is, like many biochemical systems, difficult to untangle as it is governed by a collection of complex pathways^13–16^. Upon pathway activation, complement proteins bind to the nanoparticle via thioester bonds. Bound C3bs catalyze more C3b through an amplification step, which subsequently aids in binding to other nucleophilic sites on the nanoparticle. This process triggers inflammation and recruits immune cells to the site. Furthermore, the reactions take place on a surface, as opposed to in a bulk volume. Experimental assays have thus far identified nanoparticle surface chemistry and geometry, among other factors, as an important variable influencing runaway activation^3–6,17^. While surface chemistry has been identified as an important factor, the mechanisms of this dependence are unknown.

In a recent work, we uncovered a pattern suggesting that complement spread on nanoparticle surfaces may be a critical transition. Our experimental work showed that complement activation on nanomaterial surfaces undergoes a sharp transition as a function of the mean spacing between complement attachment sites^18^. We further identified the minimal complement subnetwork responsible for this sharp transition and constructed reduced mathematical models of the collective phenomenon, including a spherical-lattice agent-based model and mean-field models, that recapitulated the critical threshold behavior observed as a function of site spacing.

Taking this finding into consideration, along with the general features of immune responses and results from spreading phenomena in other fields, we hypothesized that the complement spread may belong to the percolation universality class of critical transition. The defining feature of an immune response is the rapid deployment of immune cells given a threat and full reversibility of this deployment when the threat is resolved. There are very few classes of physical phenomena that satisfy this property. Percolation is a class of collective phase transition which would satisfy these requirements. The traditional percolation model describes thermodynamic equilibrium processes, like the Ising magnetic model. The percolation model has also been extended to dynamic processes, like epidemics, forest fires, and social opinion spreading^19–22^. More recently, studies have shown that biological systems such as neuron firing, biomolecular condensate and embryonic organization, and signaling across mitochondria and biofilm networks are examples of percolation^23–26^. However, more work needs to be done to determine whether complement spreading belongs to the percolation class, as opposed to the alternative hypothesis of a first-order Hill-type transition.

Classifying the transition undergone by complement activation on nanomaterial surfaces requires defining the susceptibility, analyzing susceptibility around the critical point with increasing system size, and analyzing critical exponents. In percolation studies, defining order parameters and susceptibilities for non-equilibrium quantities is less common than equilibrium quantities. Studies typically focus on the final cluster size or steady-state activity. In complement spread, while the final C3b coat size is of interest, we are also interested in the transition behavior of active complement proteins, a non-equilibrium quantity.

If complement spread is determined to be a critical transition, it will next be important to understand the spatial and mechanistic origin of this transition and the sensitivity of the critical behavior to environmental factors. This mechanistic model would help inform future treatment strategies and provide a framework for understanding other biochemical reaction networks on surfaces.

Here, we present a statistical-physics treatment of this non-equilibrium biochemical network to unequivocally establish that complement activation on nanomaterial surfaces is a critical transition belonging to the percolation class, rather than a first-order cooperative (Hill-type) transition. We define control and order parameters analogous to those of the canonical Ising model of percolation. We also define a susceptibility function that captures the fluctuations of C3b activation and show that this susceptibility exhibits a sharp peak at the transition. Through a systematic finite-size scaling analysis, we demonstrate that this peak diverges with increasing system size, which is a defining signature of a critical phenomenon, as distinct from a smooth, system-size-independent cooperative response. We further establish the validity and generality of this percolation picture by tracing a locus of critical points across both site spacing and intrinsic kinetic rate constants and extracting the critical exponents along this entire locus, placing the transition in a single, robust universality class. Finally, we isolate the role of molecular transport and show that the position of the critical locus is itself set by a diffusion-limited length scale, establishing that complement percolation on nanomaterial surfaces is fundamentally a transport-limited surface reaction. We close by discussing the implications of these results for nanomaterial design, immunology, and evolutionary biology.

## Methods

We combine insights from three model types, both lattice-based and mean-field, to show that complement spreading is a critical transition belonging to the percolation universality class and to illustrate the physical origins of this critical transition. While the full alternative pathway is quite complex (the full quantitative model consists of over 100 ODEs), we previously developed reduced models which agree with the full model^18,27^. Our reduced quantitative models were also validated with *in vitro* and *in vivo* experiments of complement activation by nanoparticles^18^.

### Agent-based model

To characterize complement spreading on a single-molecule level, we construct an agent-based model (ABM), described in our previous work^18^. The ABM is a quasi-2D simulation, where agents (individual C3b proteins) diffuse over a spherical surface, bind to nucleophilic sites, self-amplify, and decay.

The model contains four species: Available nucleophilic sites, active C3b-occupied sites, inactive C3b-occupied sites, and free active C3b. A free C3b will bind to an available nucleophilic site if it is within 0.075 nm. While the bound C3b is active, it autocatalyzes a new free C3b according to a catalysis probability. While the free C3b is active and unbound, it diffuses along the nanoparticle surface following a random walk with a specified diffusion speed. If the free C3b reaches an available nucleophilic site within its lifetime, it will bind to that site; otherwise, it will decay. Bound C3b agents also decay after a certain lifetime and stop producing new C3b. This process is illustrated in Fig. 1(a).

**Fig. 1.**
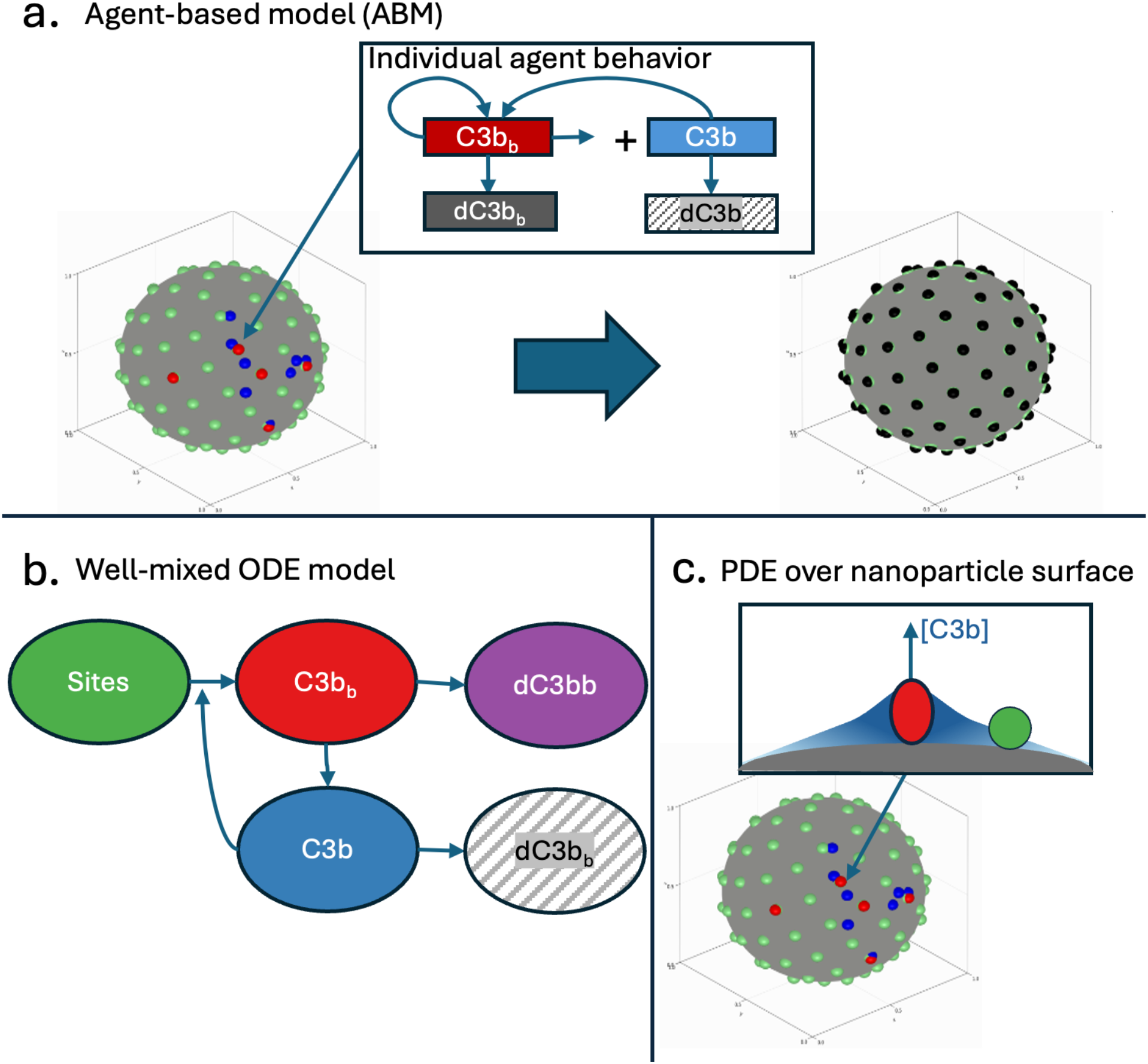
Three models represent complement physics. a.) Agent-based model (ABM) where each agent is governed by rules and life cycle and interacts with others b.) ODE model measuring the concentration of each species over time c.) Spatial, steady-state PDE measuring the concentration of C3b originating from autocatalytic site over nanoparticle surface

To vary the nucleophilic site distance on the surface, we vary the number of sites and distribute these sites on the sphere according to the Fibonacci lattice algorithm. We increase system size by increasing the radius of the nanoparticle and scaling the number of sites accordingly so that site distance stays constant.

We run each simulation for 1400 steps. Each step has time *dt* = 0.1 *τ*_*ABM*_, where *τ*_*ABM*_ is the ABM timescale. See supplementary material Table S1 for more information. Each ensemble is seeded by activating a randomly chosen site. We run 500 ensembles for each site spacing and system size. Parameters values and descriptions are given in supplementary material Table S1.

We define two order parameters for measuring complement activation, *M*_max_ and *M*_*τ*_. We define *M*_max_ as the maximum value of bound, active C3b during a single ensemble trajectory. We define *M*_*τ*_ as the activity lifetime, or the time at which activity ceases. To enable cross-comparison between nanoparticles with different site spacings and different system sizes, the reported *M*_max_ is the non-dimensional *M*_max_(*d*,size)/*N*_sites_(*d*, 1x) where *N*_sites_ is the starting number of available sites on the nanoparticle. For each order parameter, we compute the ensemble average ⟨*M*⟩ and the ensemble variance ⟨*M*^2^⟩ − ⟨*M*⟩^2^. The ensemble variance in these quantities represent the susceptibility, a classic statistical physics measurement in critical transitions, of the system, as they represent the extent to which small stochastic variations in the simulation affect the outcome^28^.

The simulations are run using Agents.jl in Julia on the Pittsburgh Supercomputing Center (PSC) Bridges-2 high-performance computing (HPC) cluster^29^. Simulating 500 ensembles for one parameter combination takes ~10 minutes on 16 CPU cores. We tested 50 site spacing values for each of four system sizes, for a total of ~32,000 CPU minutes. The order parameter statistics are computed by post-processing simulation outputs in Julia.

### Mean-field models

To characterize the full range of the critical transition and its sensitivity to the control field, we construct mean-field models of complement spreading which enable large-scale parameter sweeps.

#### Well-mixed compartment model

We first solve a time-dependent ODE model of complement spread to characterize the impact of kinetic rate parameters on complement spreading. The system is defined as

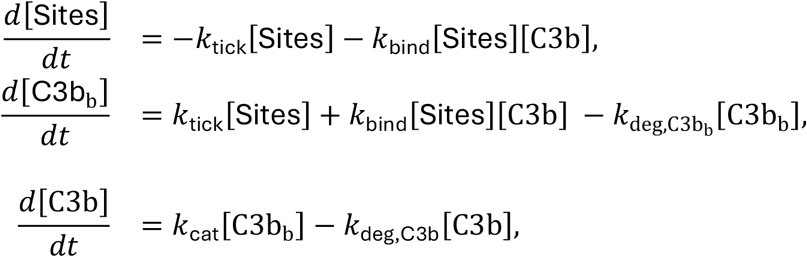

representing the evolution of concentration of available sites (Sites), active C3b-occupied sites (C3b_b_), and free active C3b (C3b). The flow between compartments is illustrated in Fig. 1(b). This model is analogous to the Targeted-Infected-Virus (TIV) model — where the empty sites correspond to Targeted cells, the free active C3b correspond to the Virus, and the active C3b-bound sites correspond to Infected cells that produce more Virus — and can be further reduced to the Susceptible-Infected-Recovered (SIR) model of disease spread^18,30^.

Our previous work determined these functions as the minimal complement subnetwork responsible for the sharp transition of complement activation. *k*_tick_ represents the rate of spontaneous tickover, or the attachment of nascent fluid phase C3b to the nanoparticle. *k*_bind_ is the attachment rate of C3b to nanoparticle site. *k*_deg,C3b_ and 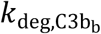 are the inactivation rates of C3b (free and bound, respectively) via the attachment of C3b to C3bBb. *k*_cat_ is the catalysis rate of C3b from a bound C3b. More details on the derivation of these reduced parameters are given in supplementary material Table S2.

The simulation is initialized with *k*_tick_ = 0 and a small initial concentration of C3b to simulate an initial tickover. We control the site distance by varying the number of sites per nanoparticle. To simulate a given site distance, we set the initial concentration [Sites] equal to the number of sites per nanoparticle times the concentration of nanoparticles. To study the sensitivity of the curves to kinetic rate constants, we repeat the same simulations by varying the values used for *k*_cat_, *k*_deg_, and *k*_bind_.

We compute *M*_max_ from the maximum value that [C3b] reaches during the simulation. Here, *M*_max_ is the dimensionless 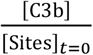, where [Sites]_*t* = 0_ is the initial concentration of available sites. We compute *M*_*τ*_ as the time when activity ceases.

These equations are solved in Julia using the DifferentialEquations.jl package with an adaptive Runge–Kutta scheme (AutoTsit5/Rosenbrock23) over the 0–600 s integration^31^. The time series plots shown are a dt=0.1s interpolation from 0-30s.

A single system can be solved in 2e-4 s. To create the order parameter versus site spacing curve, we solve the system at 246 site distance points, taking about 5e-2 s. To test sensitivity to kinetic rates, we sweep 50 values for each kinetic rate constant for a total of about 1e-2 s. To generate each of the three sets of 2D heatmaps, one set per rate constant, we run 500 values for *d* and 500 values for the rate constant. This resulted in 750k total equation solves, which takes just over 2 minutes.

We fit critical exponents to the *M* versus θ plots, where *M* is the order parameter (*M*_max_ or *M*_τ_) and *θ* is the control parameter (either *d*_*c*_ or one of the kinetic rate constants).

We compute two critical exponents:

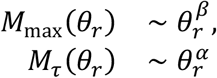

where

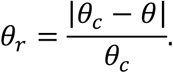

*θ*_*r*_ is the reduced control parameter based on the relative distance to critical point. Fitting details are contained in supplementary material Text S1.

#### Spatial model

We next solve a steady-state reaction-diffusion partial differential equation (PDE) on a 3D shell over a spherical nanoparticle surface to characterize the spatial origin of the critical percolation transition. The PDE is:

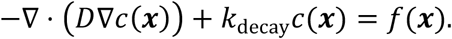

Where *c*(***x***) is the concentration profile of C3b, ***x*** is the 3D position vector in the shell, *D* is the diffusion coefficient of C3b, *k*_decay_ is the surface C3b decay rate derived from the experimentally observed half-life, and *f*(***x***) is a Gaussian point source of C3b located at the pole of the hemisphere, defined by

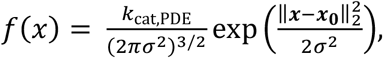

where ***x***_**0**_ is the 3D vector representing a point above the north pole of the nanoparticle and ‖⋅‖_2_ denotes the L2 norm so that 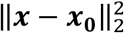 is the squared Euclidean distance of point ***x*** from ***x***_**0**_, *k*_cat,PDE_ is the C3b catalysis rate from that source point, and *σ* is the spread of the Gaussian representation of the point source. This point source represents a single active C3b-coated site, autocatalyzing C3b which diffuses across the surface. This spatial arrangement is shown in Fig. 1(c).

The computed 3d profile *c*(***x***) is then radially averaged over every point at a given angle *θ* within a thin boundary layer above the shell surface as follows:

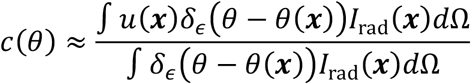

where Ω represents the 3D shell domain, *I*_rad_(***x***) is an indicator function for points ***x*** within the thin layer above the shell surface, and *δ*_*ϵ*_ is a smoothed delta function for points ***x*** near angle *θ*. Results are presented as a function of surface arc length *s* = *θ* ∗ *R* nm, where *R* is the radius of the sphere.

We use zero-flux Neumann boundary condition on the sphere surface and Dirichlet zero-value boundary conditions on the outer boundary. This model is solved in Julia using the finite element method in the Gridap.jl package, completing in about 20 minutes on one CPU^32,33^. To produce each of the three 2D parameter sensitivity sweeps, we tested seven by seven grids of points ranging from 0.1-10x the base parameter value for each parameter. This resulted in ~3000 CPU minutes, completed in parallel on PSC Bridges-2 HPC cluster^29^. The base parameter values for *k*_cat,PDE_, *k*_decay_, and *D* are given in supplementary material Table S3.

## Results

### Establishing complement spreading as belonging to the percolation universality class of critical transition

In this section, we first demonstrate that complement spreading on nanoparticles is a critical transition by showing it satisfies properties of critical transitions. Our novel, lattice-based ABM model of complement adhering to nanoparticle surfaces uniquely allows us to interrogate these properties in our system for two main reasons: 1.) As a stochastic model, the model generates ensembles of possible outcomes, allowing us to analyze probability distributions and fluctuations of the order parameters (Fig. 2) and 2.) As a spatially explicit model, we are able to investigate the effect of system size.

**Fig. 2.**
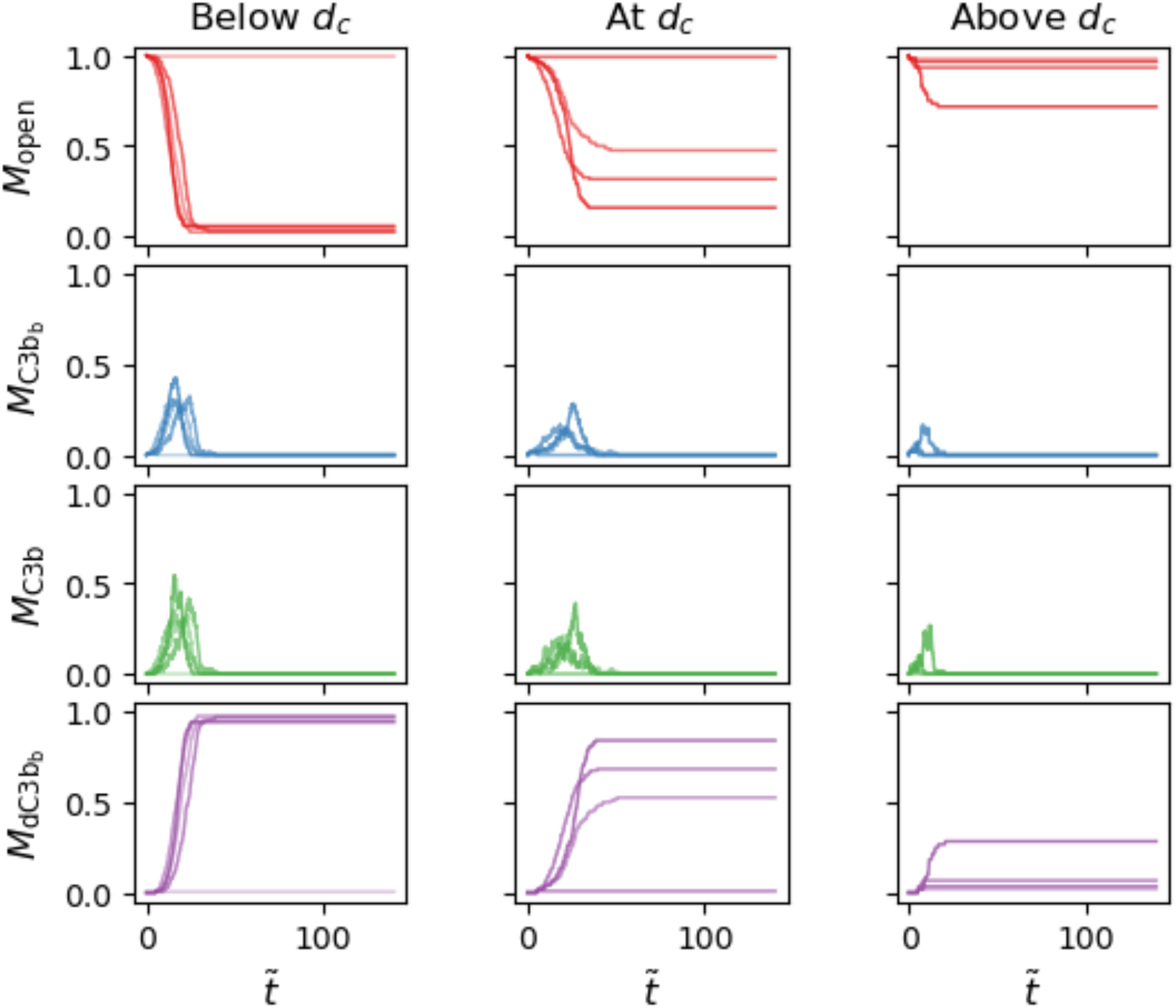
Complement ABM produces stochastic trajectories for a.) open sites (M_open_) b.) Active C3b-bound sites 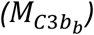 c.) Free, active C3b (M_C3b_) d.) Decayed C3b-bound sites 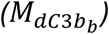. Each line represents the trajectory of an individual ensemble. Each column represents a different site spacing.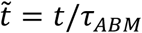, a non-dimensional time with respect to the simulation time scale τ_ABM_.

To characterize the percolation transition, we report the ensemble mean and variance of *M*_max_ and *M*_τ_ (defined in Methods) and show that these follow the transition and scaling behavior expected from critical transitions. The ensemble means of each observable versus site spacing are shown in Figs. 3(a) and (b), and the variances are shown in Figs. 3(c) and (d). ⟨*M*_max_⟩ shows a sharp power-law like transition with respect to site spacing, suggesting a critical transition between the regime where C3b dies out before it spreads and the regime where C3b produces runaway events^19,20,28^. ⟨*M*_*τ*_⟩, on the other hand, shows a divergence around the critical transition point, capturing a critical slowing down^20^. Both Var(*M*_max_) and Var(*M*_*τ*_), the susceptibility metrics in our model, exhibit the expected divergence around the critical point^20,28^.

**Fig. 3.**
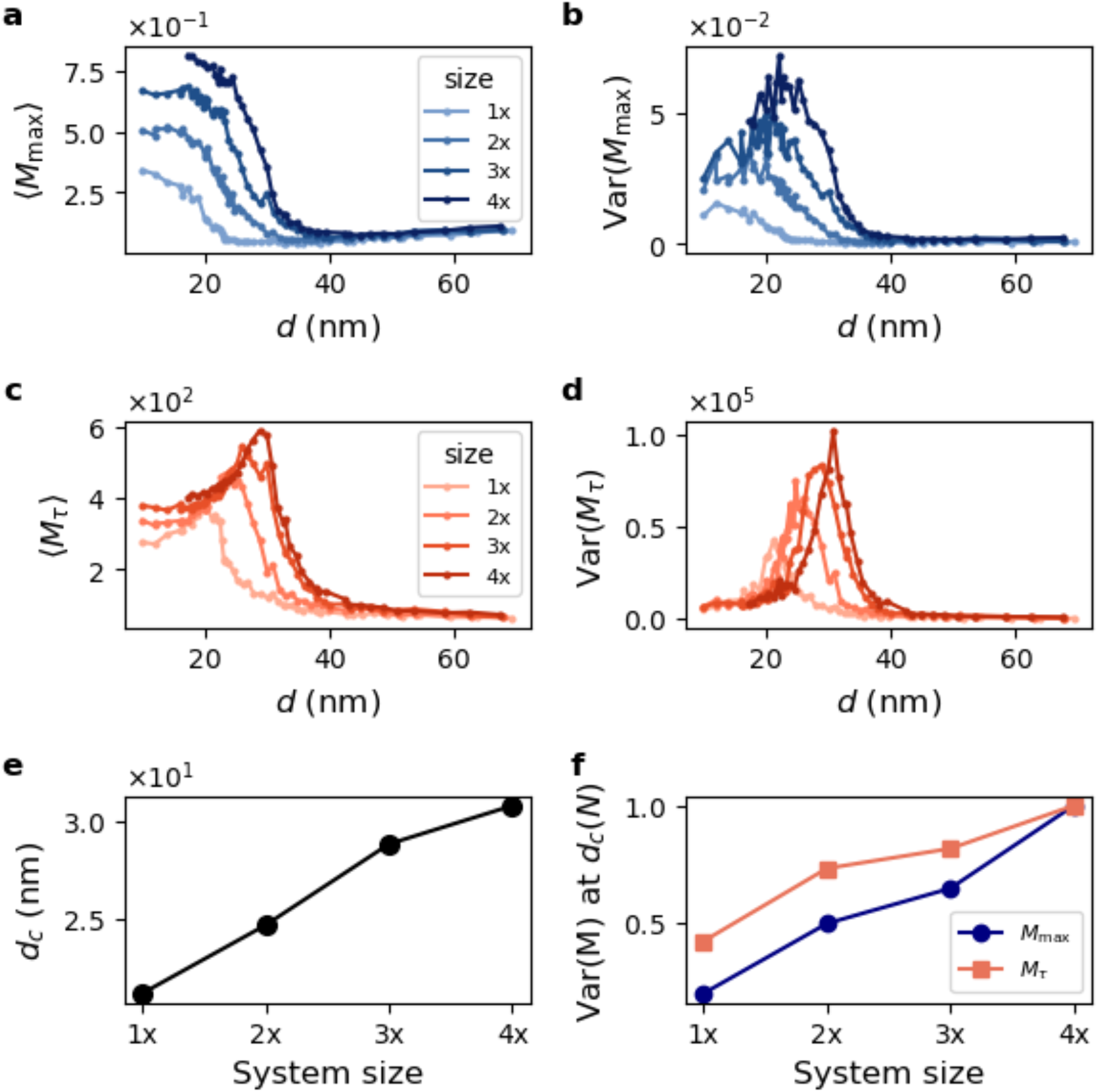
Complement ABM shows hallmarks of critical transitions across different system sizes N. a.) Maximum activity versus site spacing. b.) Variance in maximum activity versus site spacing c.) Lifetime versus site spacing d.) Variance in lifetime versus site spacing e.) d_c_ versus system size, with d_c_ estimated from the variance peak location f.) Variance peak height versus system size, normalized by the value at 4x.

To illustrate the susceptibility in our model, the supplementary material Movie M1 presents two spatiotemporal trajectories for each of three ABM site spacings. For site spacings near and below the critical point, there are two possible outcomes: 1.) C3b dies out before spreading or 2.) C3b covers most of the nanoparticle. These disparate outcomes indicate high susceptibility. For site spacings above the critical point, C3b always dies out quickly before spreading. The lifetime of the high activity ensemble at the critical point is greater than that of the one below the critical point, illustrating the lifetime divergence. Consequently, there are more runaway ensembles for site spacing below the critical point; thus, while the system is still susceptible to fluctuations, the susceptibility decreases below the critical point, illustrating the susceptibility divergence at the critical point. Note that the sphere surface in the movie is transparent so that all attachment sites are visible.

This shift and divergence in behavior can be further illustrated by plotting histograms of *M*_max_ and *M*_τ_ with changing *d*, seen in Figs. S2-3. At *d* >> *d*_*c*_, most values land on the low end of the histogram, representing trajectories dying out almost immediately. As *d* → *d*_*c*_, the distribution spreads out from left to right, representing the shift to runaway trajectories, with ensemble outcomes governed by stochasticity. At *d* < *d*_*c*_, most values land on the high end of the histogram, representing the runaway activation regime.

We also observe crucial system size scaling behavior for critical transitions^20,28^. As system size increases, the critical point increases and shows evidence of convergence towards a true, infinite-size system critical point value (Fig. 3(e)). In addition, the divergent susceptibility-related peaks increase and sharpen with increasing system size (Fig. 3(b)-(d) and (f)). Thus, even though a first-order Hill function could also fit the activation versus site distance curve (Fig. S4), the fluctuation and system-size scaling results show that it must be a critical transition.

While we highlight activity-related quantities in Fig. 3, we note that the final C3b-consumed cluster size, analogous to the canonical thermodynamic percolation order parameter, also follows a percolation transition characterized by divergence, shifting histogram distributions, and system-size scaling^20^. These results are shown Figs. S1 and S5. This behavior matches that of experimental observations, further validating our model^18^.

### Establishing critical locus for complement spreading percolation

In this section, we use the mean-field models to characterize a locus of critical points and exponents as a function of the kinetic rate parameters in addition to the site distance, *d*_*c*_.

We first compare the mean-field model results to those of the ABM and show that it captures the same percolation transition. In contrast to the ABM’s ensemble of stochastic trajectories, each parameter set in the mean-field produces a smooth deterministic trajectory, which is also observed by plotting the ABM ensemble average trajectory ⟨*M*_*c*3*b*_(*t*)⟩ seen in Fig. S6. To show the mean-field model captures the same percolation transition as the ABM, we plot the same two order parameters, *M*_max_ and *M*_*τ*_, as a function of site-spacing. The same critical transition of these two order parameters observed in the ABM is observed in the mean-field model, with a power-law transition in *M*_max_ and divergence in *M*_*τ*_, shown in Figs. 5(a) and (d).

**Fig. 4.**
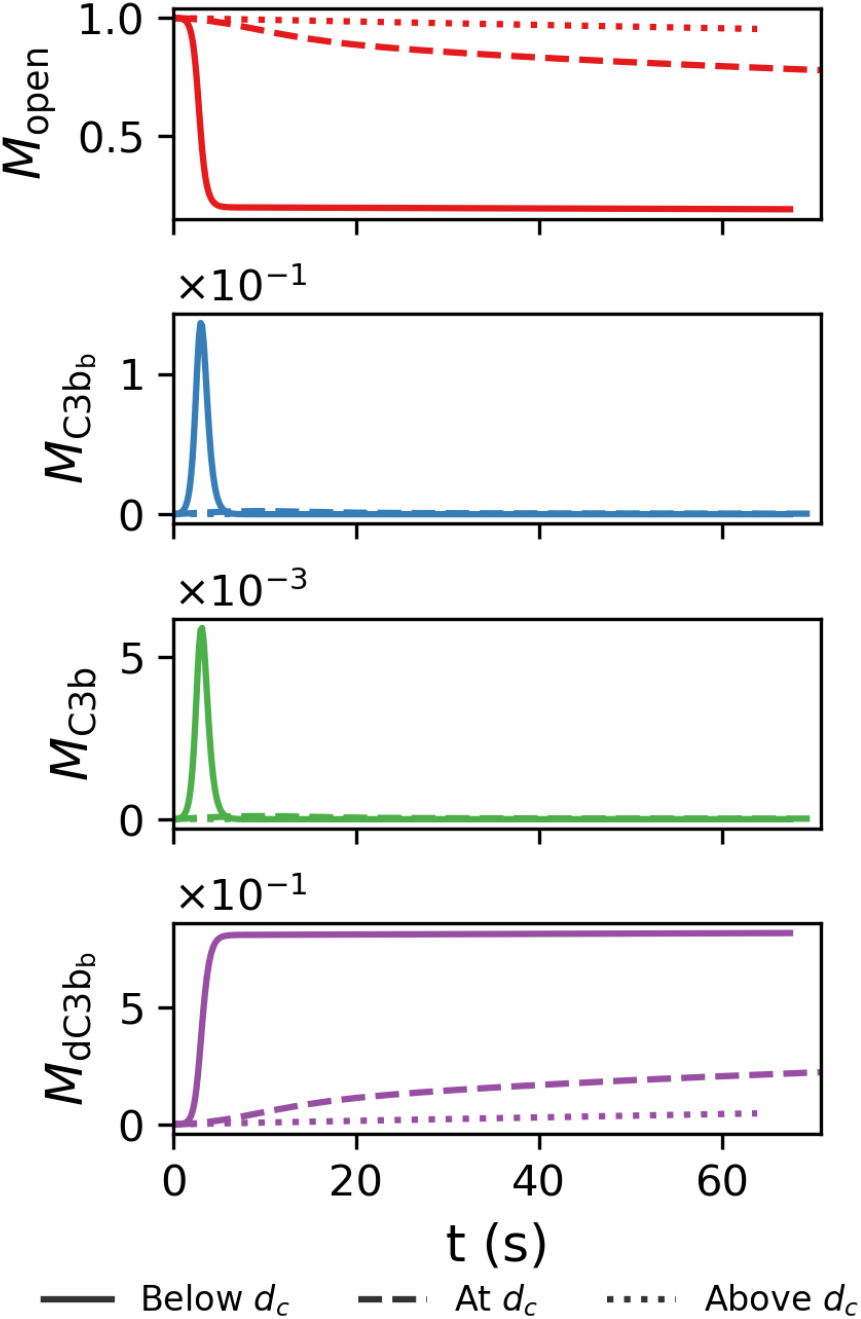
Complement TIV model trajectories for a.) open sites (M_open_) b.) Active C3b-bound sites 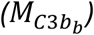 c.) Free C3b (M_C3b_) d.) Inactive C3b-bound sites 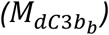. Each line represents the trajectory for a different site spacing.

**Fig. 5.**
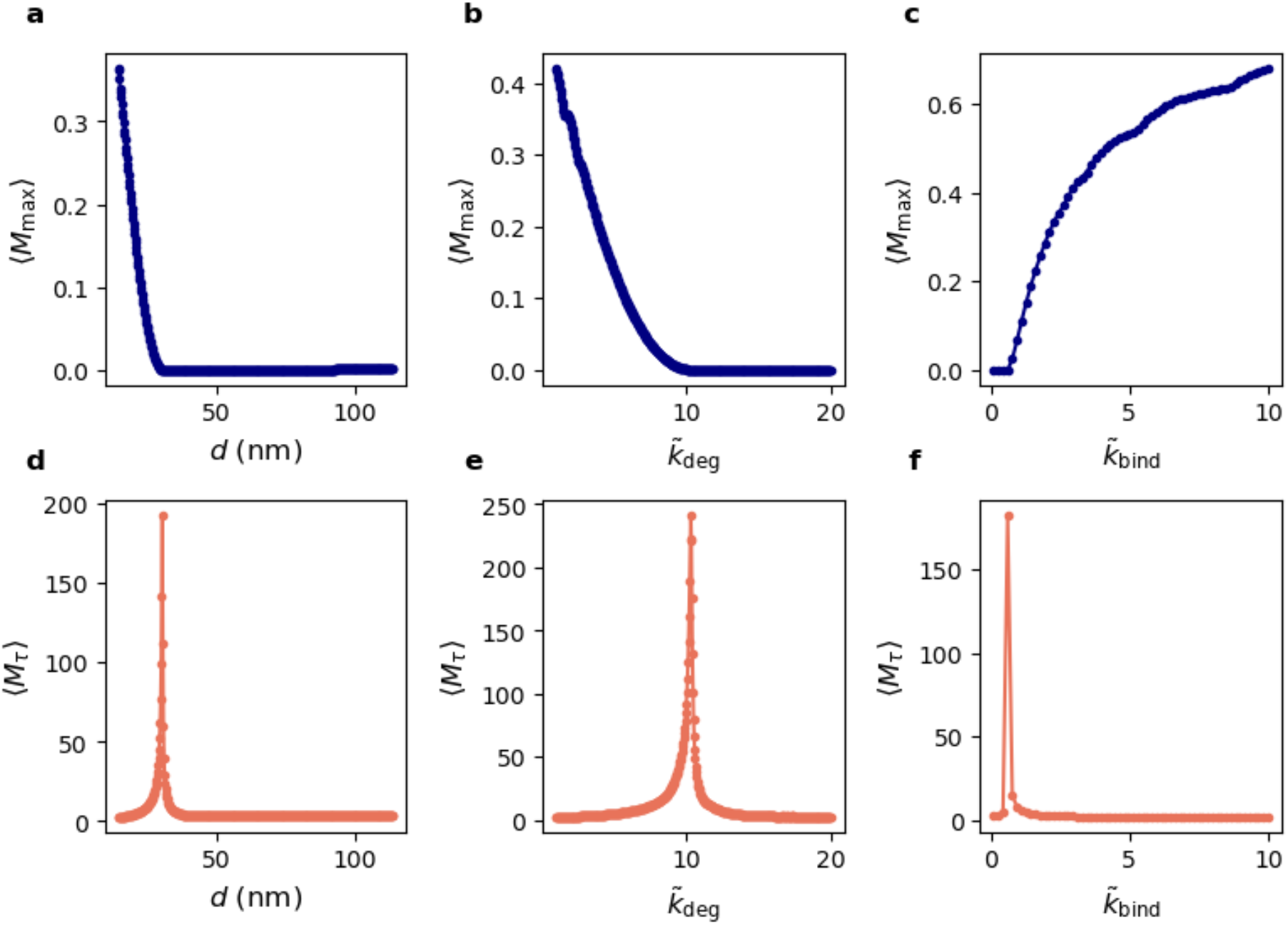
Well-mixed mean-field model of complement demonstrates hallmarks of critical transitions as a function of both site spacing (first column) and kinetic rate scaling (second and third columns). The top row shows maximum activity M_*max*_ versus control parameters. The bottom row shows lifetime M_τ_ versus control parameters. *Parameter values are presented with respect to their baseline values:* 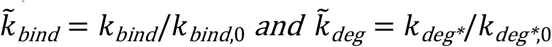 *(the * denotes either C3b or C3b*_*b*_, *as the two degradation rates are scaled equally and simultaneously). k*_*bind*,0_ and 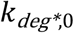 refer to the baseline TIV parameters given in supplementary material Table S2.

We next test how the system responds to perturbations to kinetic rate constants. Since *k*_deg,C3b_is a constant multiple of 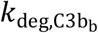, we simultaneously scale both degradation rates so that this ratio is maintained. We find that critical transitions in *M*_*max*_ *and M*_*τ*_ are additionally observed as a function of each kinetic rate, shown in Figs. 5(b)-(c),5(e)-(f), and S7. The presence of these additional critical transitions suggests the existence of a critical regime dependent on both site distance and kinetic rate parameters, analogous to the canonical Ising model dependence on the coupling constant and temperature. This finding leads us to establish a locus of critical behavior by performing 2D parameter sweeps across different combinations of site distances and kinetic rate parameters, with results shown in Figs. 6(a)-(b). The plots in Fig. 5 represent individual horizontal and vertical slices of these phase diagrams in Fig. 6. The thin line in Fig. 6b represents the location of the critical locus. Critical transition plots and heatmaps for all reaction rate constants and order parameters can be found in Figs. S7-8.

**Fig. 6.**
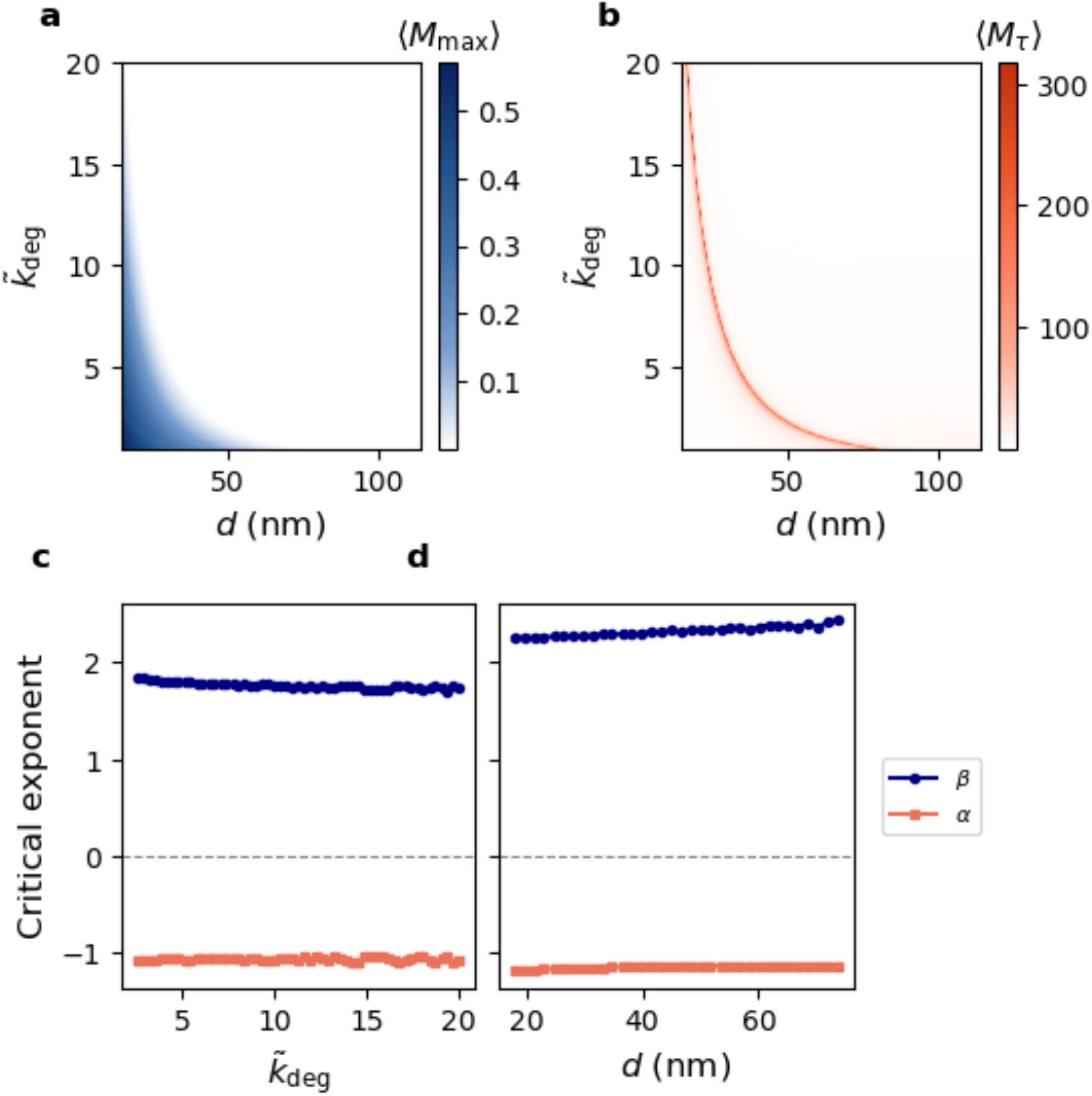
Complement model generates critical locus. a-b) Heatmaps of max active (a) and lifetime (b) as a function of 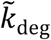 and d. c-d) Critical exponents computed across the locus. c.) critical exponents for *M*_*max*_ and *M*_*τ*_ versus site spacing d as a function of kinetic rate scaling 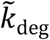. d.) critical exponent for *M*_max_ and *M*_*τ*_ versus kinetic rate scaling 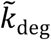 as a function of site spacing d. 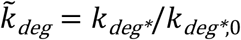 *(the * denotes either C3b or C3b*_*b*_, *as the two degradation rates are scaled equally and simultaneously)*, where 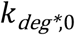 refers to the baseline TIV parameter given in supplementary material Table S2.

We next characterize the critical exponents across the locus following the fitting method described in Methods. We find that each exponent has a distinct value which remains consistent across the locus. As shown in Fig. 6(c) and (d), the empirical fits give *β* = 2 for the peak activity exponent and *α* = −1 for the lifetime exponent. Both critical exponents can be derived directly from the mean-field reaction network near the critical point (supplementary material Text S2) giving *α* = −1 (supplementary material Text S2.2) and *β* = 2 (supplementary material Text S2.3) in exact agreement with the empirical fits. The lifetime exponent *α* additionally matches the mean-field dynamic exponent of the general epidemic process, while the peak activity exponent has not previously been derived from mean-field^19^.

### Establishing transport origin of percolation

In the previous section we showed that critical behavior was dependent on kinetic rate constants. Here, we reveal the spatial constraints underlying this effect. The reaction-diffusion model (described in Methods) captures the requirement that nascent C3b must diffuse across the nanomaterial surface to a nearby attachment site to bind and autocatalyze before it decays. By solving the reaction-diffusion PDE model with varying catalytic rate (*k*_cat_), decay rate (*k*_decay_), and diffusion coefficient (*D*), we show that complement percolation is transport-limited. Solved profiles are shown in Fig. S9. When the inter-site spacing exceeds a critical distance, most nascent C3b hydrolyzes prior to binding, suppressing amplification; when spacing lies below this threshold, C3b can propagate across the surface and initiate the amplification loop. If the diffusion length increases, the critical distance at which this transition occurs increases. The heatmaps in Fig. 7 illustrate the impact of the physical parameters on the diffusion length. We compute the diffusion length *λ* by fitting the exponential function

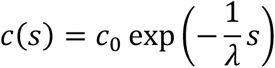

to the numerically solved profile. The heatmaps are colored by *λ*. As *D* increases and *k*_deg_ decreases, the diffusion length increases. While *λ* remains constant with increasing *k*_cat_, *u*_*o*_ increases as *k*_cat_ increases (shown in Fig. S9 inset). Thus *c*(*λ*) increases with increasing *k*_cat_, also increasing the critical distance. These findings agree with the critical point increasing with *k*_cat_ and *k*_bind_ in Fig. S8 and decreasing with *k*_deg_ in Fig. 6. This analysis agrees with our previous finding that changing the diffusion speed in the ABM shifts the critical behavior^18^.

**Fig 7.**
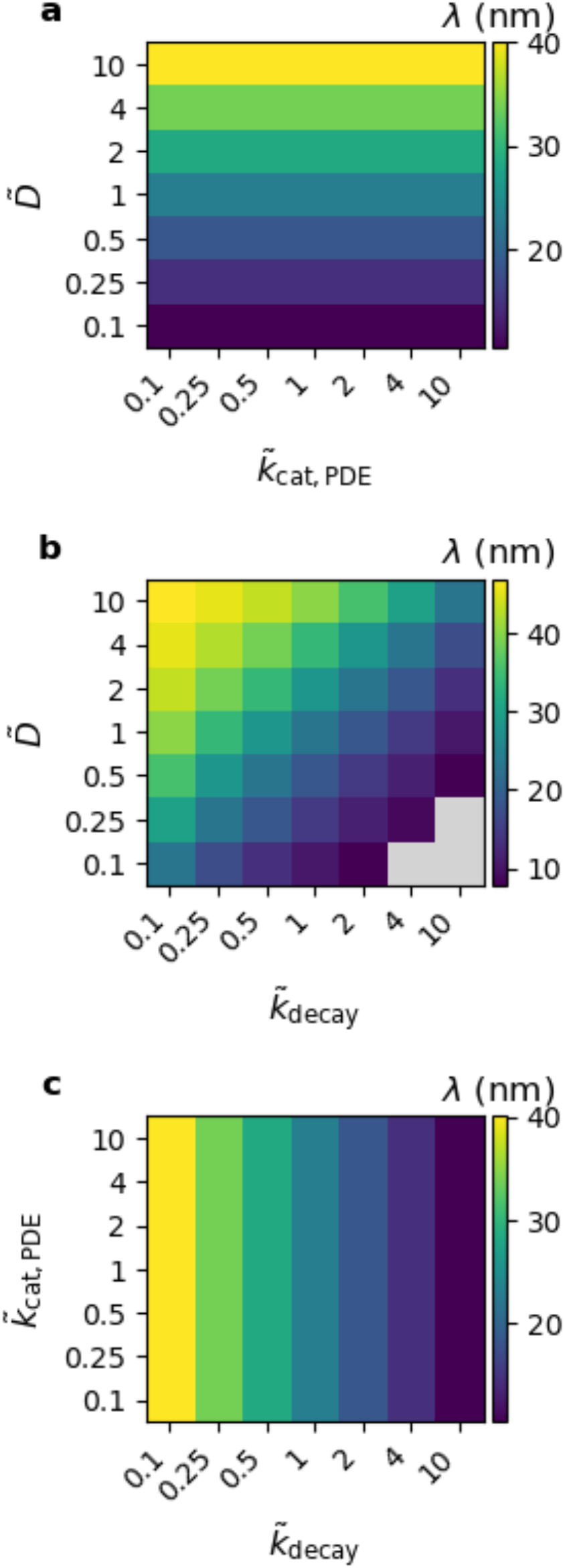
*Heatmaps of parameter combinations in the PDE a.) Diffusion coefficient* 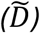 *versus source catalytic rate* 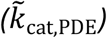 *b*.*) Diffusion coefficient* 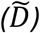 *versus decay rate* 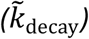 *c*.*) Source catalytic rate* 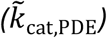 *versus decay rate* 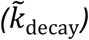. *Parameter values are presented with respect to their baseline values:* 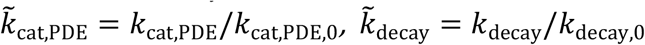, *and* 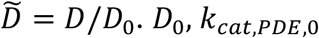, and *k*_decay,0_ refer to the baseline PDE parameters given in supplementary material Table S3.

We also previously found that increasing the tickover rate shifts the critical point. The tickover rate represents the frequency at which a random nucleophilic site is seeded with nascent, fluid-phase C3b^18^. In the results in the present study, we randomly seeded just once at the beginning of the simulation. While the tickover rate does not influence connectivity between nodes, it can increase the probability of a nanoparticle-covering cluster forming as the union between two smaller clusters seeded by two tickover events. Thus, increasing this rate also shifts the critical point.

## Discussion

Understanding the nature and physical origin of the complement activation transition is a key challenge in designing safe and effective therapeutic nanomaterials and understanding complex immune-related disease processes. In this study, we apply statistical-mechanics tools on hierarchical models to characterize a phase transition in a non-equilibrium biochemical surface reaction. Analogous to canonical magnetic Ising model percolation where spin correlation is a function of both the coupling constant *J* and temperature *T*, our model of complement activation is a function of both the nucleophilic site coupling via site distance (*d*_*c*_) and the kinetic rates (representing “hot” versus “cold” immune environments).

Using our model, we demonstrate divergence in the susceptibility function through system size scaling analysis and delineate the locus of critical points and exponents.

First, we use the ABM to show that the complement spreading transition as a function of surface site spacing is a critical transition belonging to the percolation universality class, as opposed to a first-order transition. By representing the spatial and stochastic elements of complement spread, the model reveals that complement spreading exhibits hallmarks of critical transitions like fluctuation divergence, system-size scaling, and critical slowing down.

While susceptibility in the Ising model refers to equilibrium thermodynamic fluctuations, our model’s susceptibility refers to how much the stochastic variance in each ABM step affect the overall trajectory. Below the critical point, nearly all runs die out before propagating, regardless of stochasticity in the steps. Above the critical point, nearly all runs runaway regardless of stochasticity in the steps. At the critical point, trajectories may either die out or propagate, varying greatly due to model stochasticity. The maximum activity and activity lifetime both represent different aspects of trajectory outcome. Thus, studying their variance represents the susceptibility of our system. The variance of both quantities diverges at the critical point, supporting that our system is a critical phenomenon.

The activity lifetime itself also experiences a divergence. Below the critical point, the system quickly dies out. Above the critical point, the system quickly consumes the whole system. At the critical point, the site spacing is at the edge of where C3b can diffuse before decaying, so the activation is able to runaway but takes a long time to do so. This dynamic timescale diverges for the same reason that susceptibility diverges. Thus, we can observe the divergent behavior at the critical point even in the mean-field model which does not capture stochastic fluctuations.

Using the well-mixed mean-field and spatial mean-field models (ODE and PDEs), we show that the critical transition of complement activity is also governed by kinetic rates and transport processes. We determined the existence of a critical locus spanning site spacing and physical rate constants with a set of consistent critical exponents across the locus. Our models are validated by our previous experimental data and furthermore enable the study of stochastic, temporal dynamics beyond what can be observed experimentally. Our final cluster size results agree with models of spreading process percolation in other fields, like epidemics, forest fires, and social opinion spreading^19–22^. In contrast to many of these models, we focus here on the peak number of active agents during the complement spreading process. We found only one other study of dynamic percolation that reports this peak activity^34^. However, this measure is important to consider in the context of immune reactions. Active C3b autocatalysis releases anaphylatoxins. Anaphylatoxins in high amounts will both increase the number of macrophages recruited to the site, limiting biodistribution of the nanoparticle via phagocytosis, and increase the risk of shock. Observing this peak activity — and furthermore the fluctuations, system-size scaling, and lifetime of the activity — is difficult experimentally. Our simulation models enable tracking these latent quantities. Our models also enable studying the impacts of large sweeps of rate constants, which otherwise would require mapping serum composition to rate constants and precisely modulating the serum composition and animal model immune systems accordingly.

Our work contributes to the growing thesis that 1. biology has evolved to operate at criticality^35^, and that 2. onsets of unwanted reactions and disease processes are critical transitions that can be detected and managed^36,37^. For instance, studies have shown that phenomena like neuron firing, biomolecular condensate and embryonic organization, and signaling across mitochondria and biofilm networks are examples of percolation^23–26^. Furthermore, hallmarks of critical transitions have been shown to be associated with disease transitions in epilepsy, depression, heart disease, neurodegenerative disease, and immune activation after cardiac surgery, among others^36,38^.

The immune system residing at criticality, and that it is a critical transition as opposed to a first-order transition, has many interesting implications for biology. For example, perhaps it is beneficial for the system to reside near criticality so that shifting to an activated, pathogen-attacking state has a low energy requirement. This logic has been applied to explaining why neural activation exists at criticality: neurons fire just enough to enable cognitive processes but not too much to lead to seizures^35^. Furthermore, the reversibility of a critical transition enables restoration of the less-activated state when it is no longer needed; however, if one of the nodes is perturbed too much (e.g. due to a mutation or other influence), then the typical controls may be insufficient. In contrast, first-order transitions, which may represent chronic disease transitions, are more difficult to reverse after the tipping point due to bifurcations.

Complement modulation on nanoscale surfaces is also integral to how host cells, pathogens, and cancer cells evade the immune system. Host cells avoid immune recognition because their membranes present proteins that recruit complement degradation factors^8^. Cancer cells and pathogens also present these proteins, suggesting that they may have evolved to produce an optimal amount of these factors based on the critical transition point to prevent complement adhesion^8,9^.

The findings provided by our work could be used to inform design strategies for nanotherapeutics. Recent design strategies to reduce complement activation have included conjugating C3b inactivation factors to the nanocarrier surface and injecting of immune modulation compounds alongside the nanomedicine^39–42^. Our model could help optimize these kinds of strategies, by showing how the changes in kinetic rates caused by these additional factors would affect the resulting complement activation. Furthermore, as experimental techniques allow precise adhesion of functional groups to nanocarrier surfaces, our model can help optimize this adhesion.

The findings provided by our work can also be used to inform therapeutic strategies for disorders triggered by complement adhesion to unwanted nanosurfaces in the body, such as in age-related macular degeneration (AMD) and Alzheimer’s Disease (AD)^10,11^. The model can help define which nodes in the complement pathway are key to target and to what extent.

In addition to complement, our hierarchical modeling approach provides a framework for generalizing to modeling other biochemical processes to study transitions in their behavior. Our model can study unobservable dynamics and identify the response function to specific, tunable, physical parameters. We could then create a machine learning surrogate model of these simulations to enable even faster characterization and inverse design of nanoparticles. Finally, combining insights from multiple model scales enables a more holistic understanding of the process.

## Supplementary Material

The supplementary material contains figures S1-S9, tables S1-S3, movie M1, and sections S1-S2. These components display additional simulation analyses (including order parameter histograms, Hill function fit, full control parameter sweeps, and PDE profiles), model parameters, exponent fitting methodology and analytical derivations, and a movie of agent-based model trajectories.

## Acknowledgments

*This work has been supported in part by the National Institutes of Health under grants NIGMS GM136259 and NCI CA250044. This work used Bridges-2 at Pittsburgh Supercomputing Center through allocation MCB200101 from the Advanced Cyberinfrastructure Coordination Ecosystem: Services & Support (ACCESS) program, which is supported by National Science Foundation grants #2138259, #2138286, #2138307, #2137603, and #2138296, and through the Penn Advances Research Computing Center’s Betty cluster*.

## Data Availability Statement

Supplementary Information is provided online. Code will be available on GitHub following publication. Data will be available in Zenodo following publication.

## Supplemental Figures

**Fig. S1.**
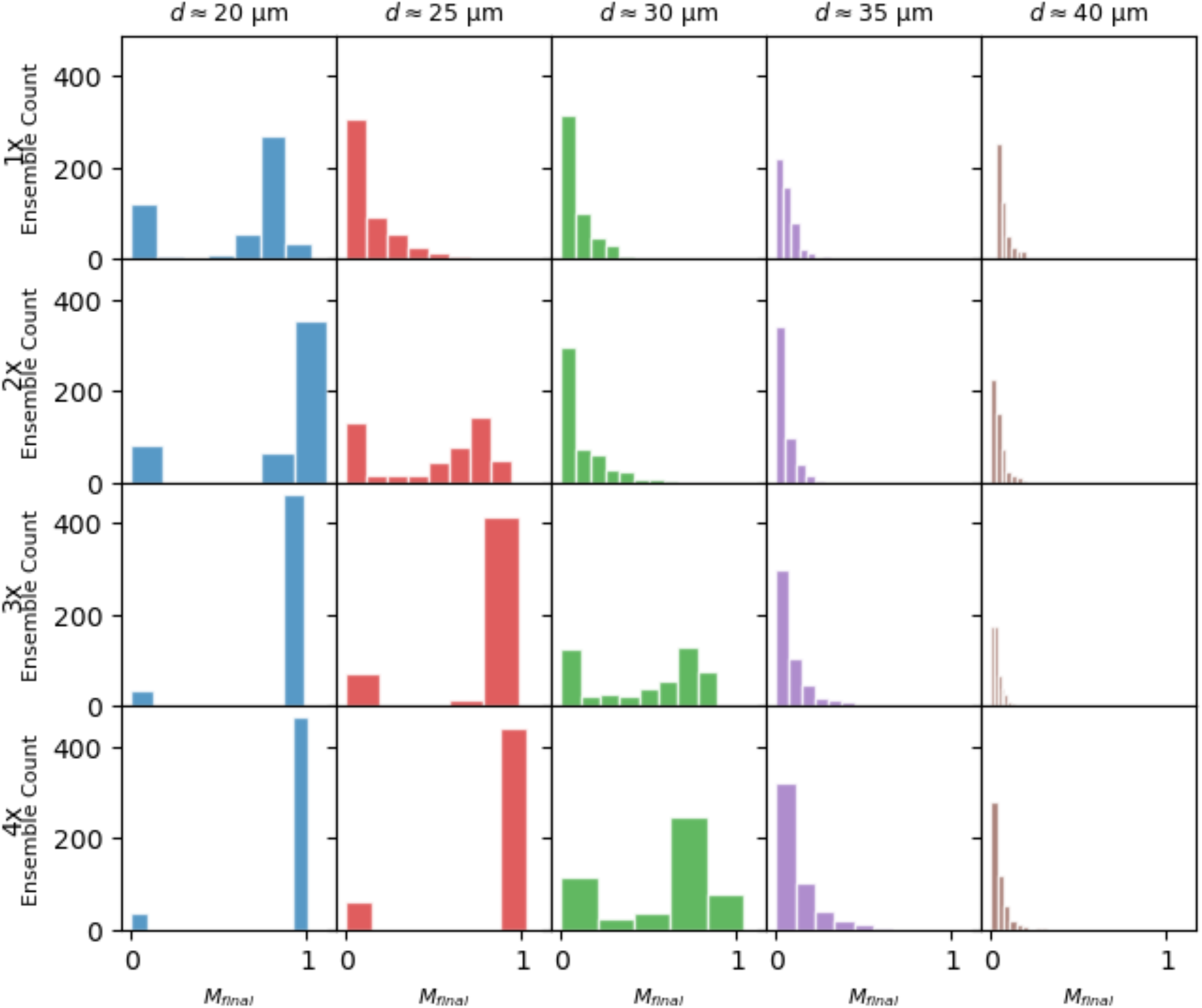
Ensemble distributions of M_fiinal_ for select site spacings with increasing system size. Each column represents a site spacing and each row represents a system size.

**Fig. S2.**
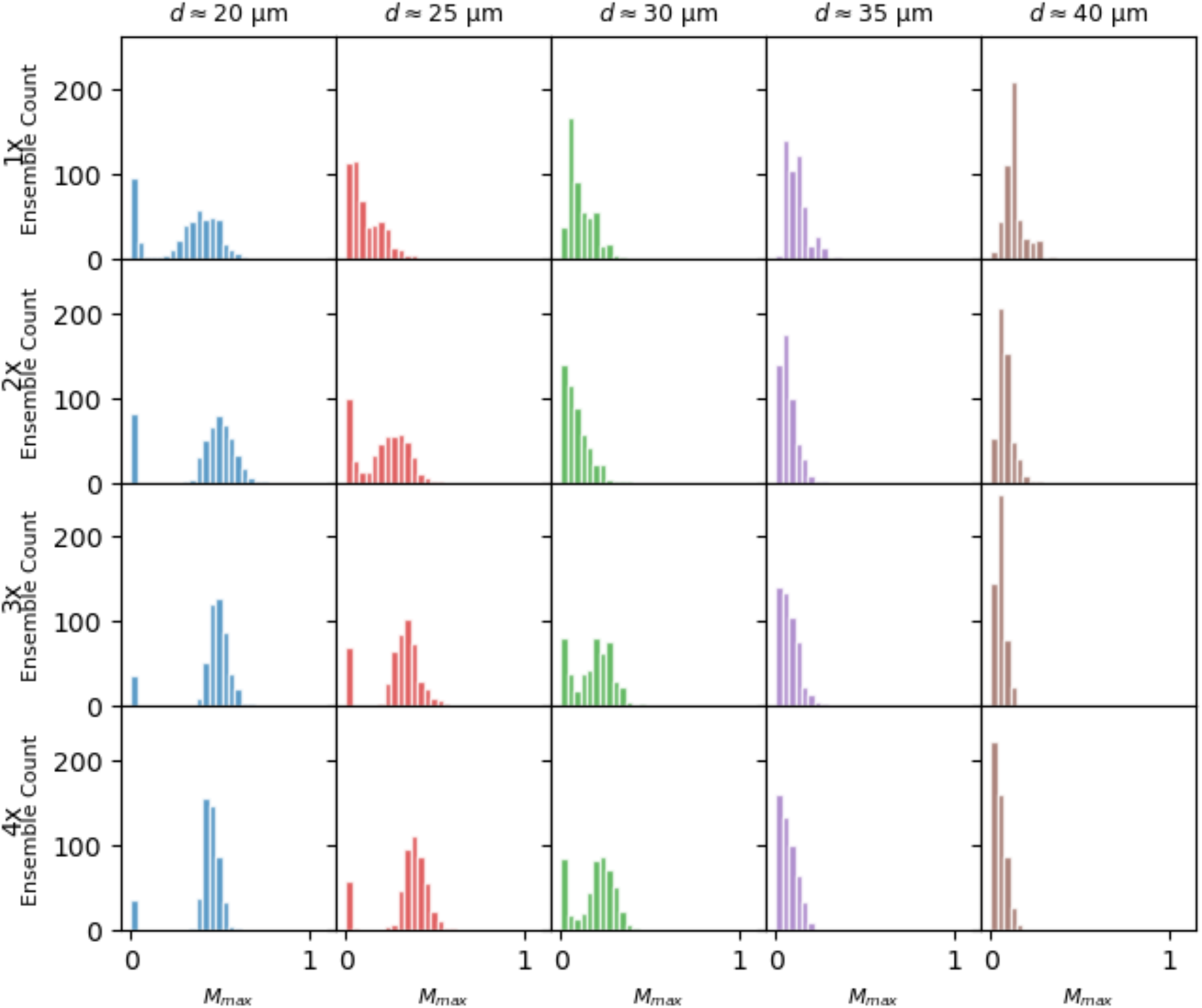
Ensemble distributions of M_max_ for select site spacings with increasing system size. Each column represents a site spacing and each row represents a system size.

**Fig. S3.**
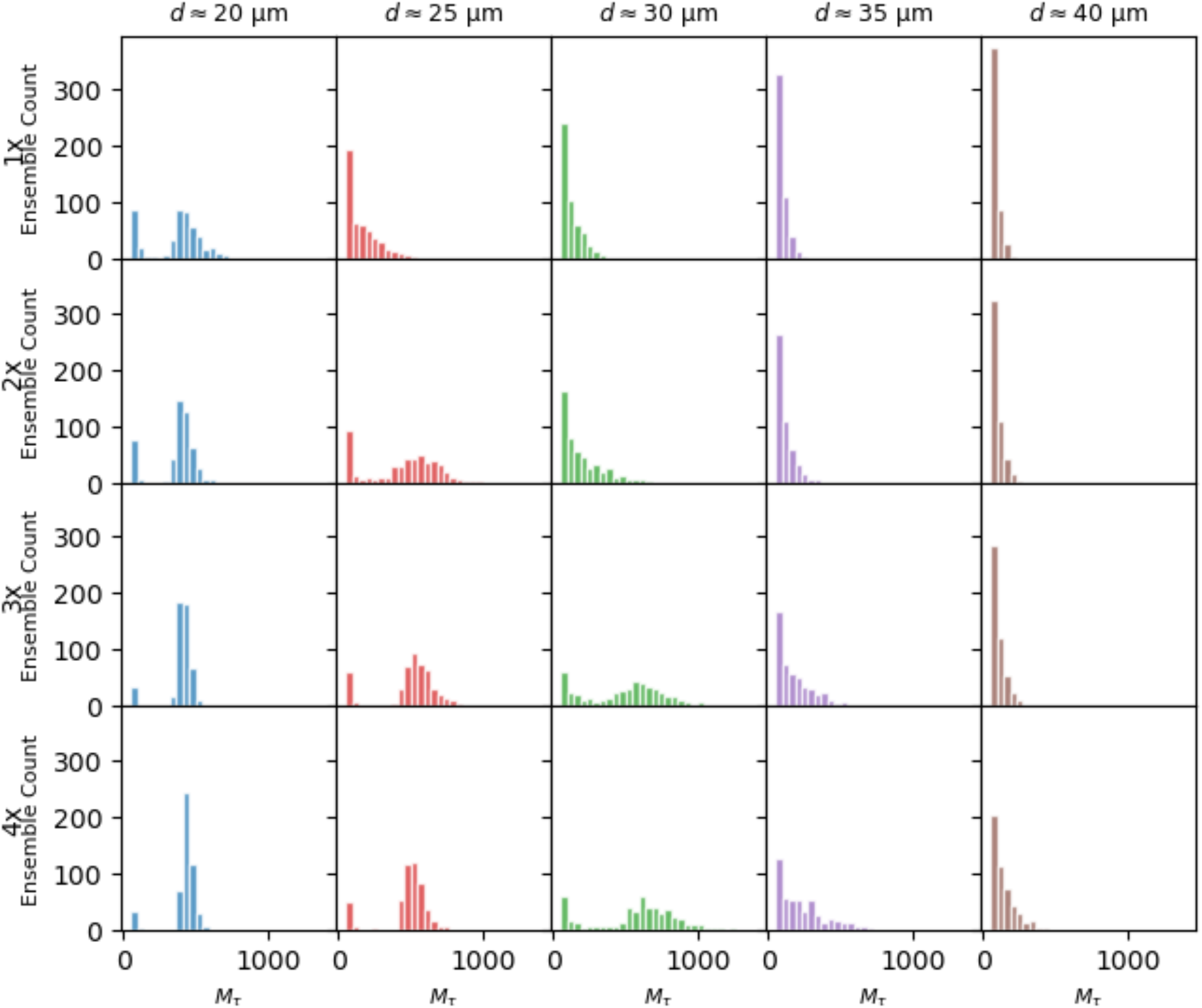
Ensemble distributions of M_τ_ for select site spacings with increasing system size. Each column represents a site spacing and each row represents a system size.

**Fig. S4.**
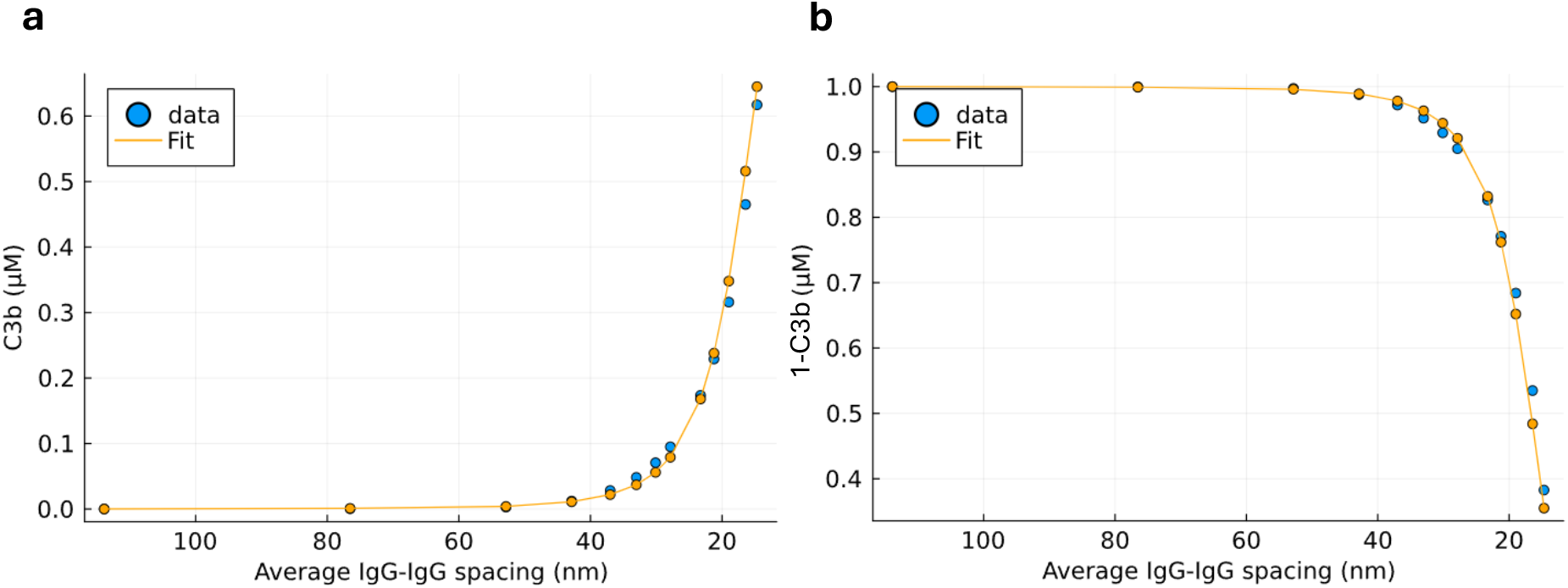
*Simulation data with Hill function fit: a.*) *C3b and b.) Consumed sites. The data is fit to the Hill function form* 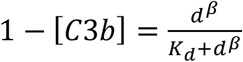. We use an ordinary least-squares fit on 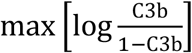 versus distance. We find *β* = 4.7, giving the fit model 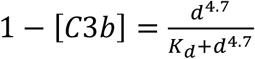.

**Fig. S5.**
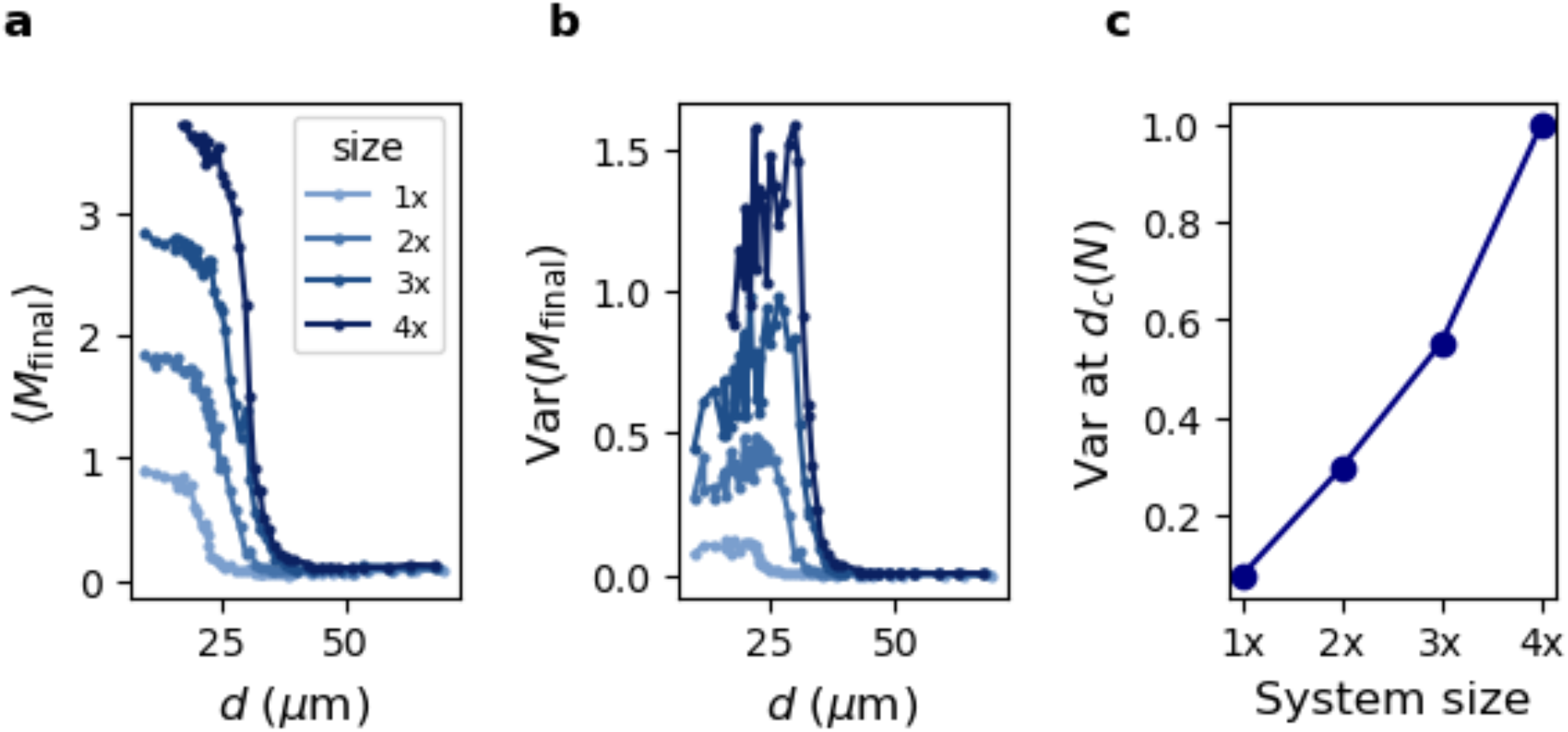
Mean, fluctuations, and system-size scaling of final occupied sites, M_fiinal_, order parameter in ABM simulations.

**Fig. S6.**
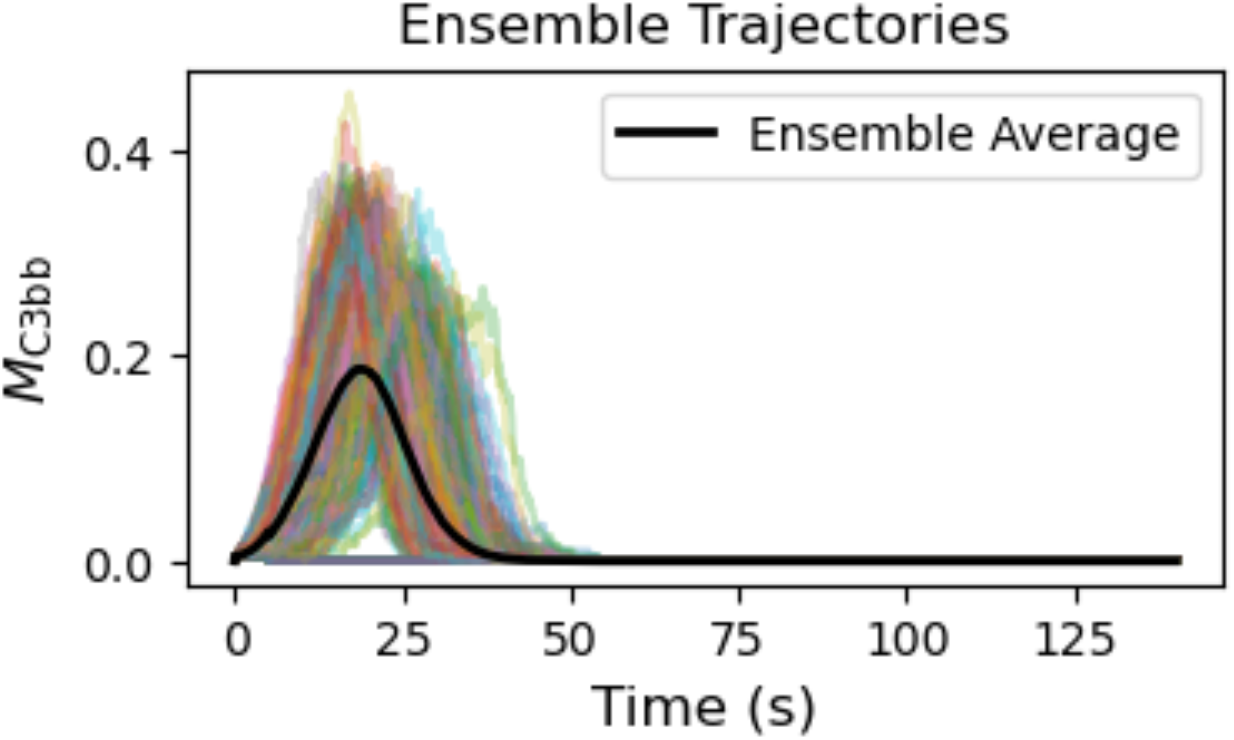
Stochastic ABM trajectories for N=500 ensembles (multi-colored lines) and the trajectory average (black line).

**Fig. S7.**
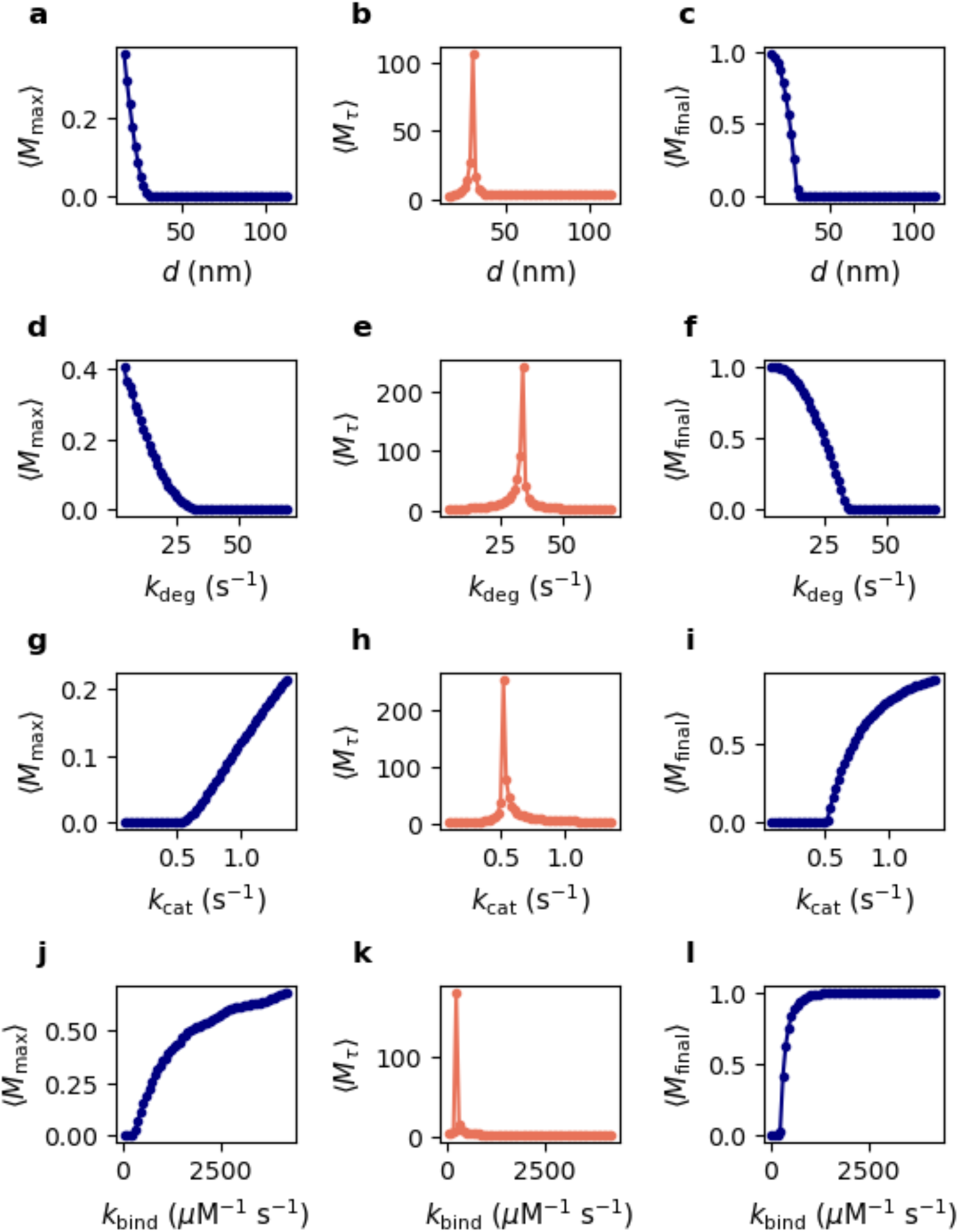
Order parameters versus site distance, *d*, and kinetic rate constants, *k*_deg_, *k*_cat_, and *k*_bind_. Each column is a different order parameter; each row is a different control variable.

**Fig. S8.**
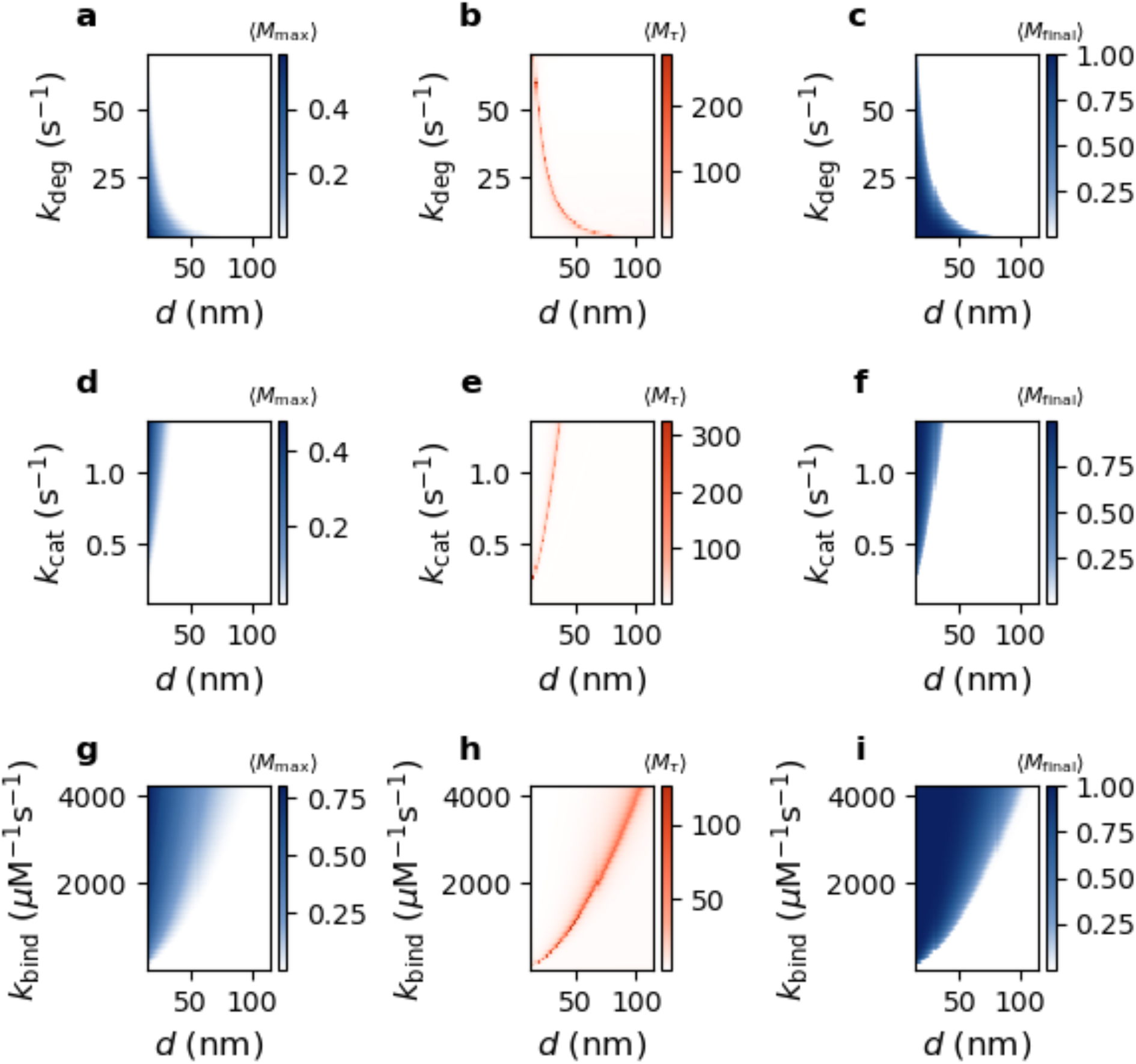
Heatmaps of order parameters as 2D function of site distance, *d*, and one of the kinetic rate constants (*k*_deg_, *k*_cat_, or *k*_bind_). Each row represents a different kinetic rate constant; each column represents a different order parameter.

**Fig. S9.**
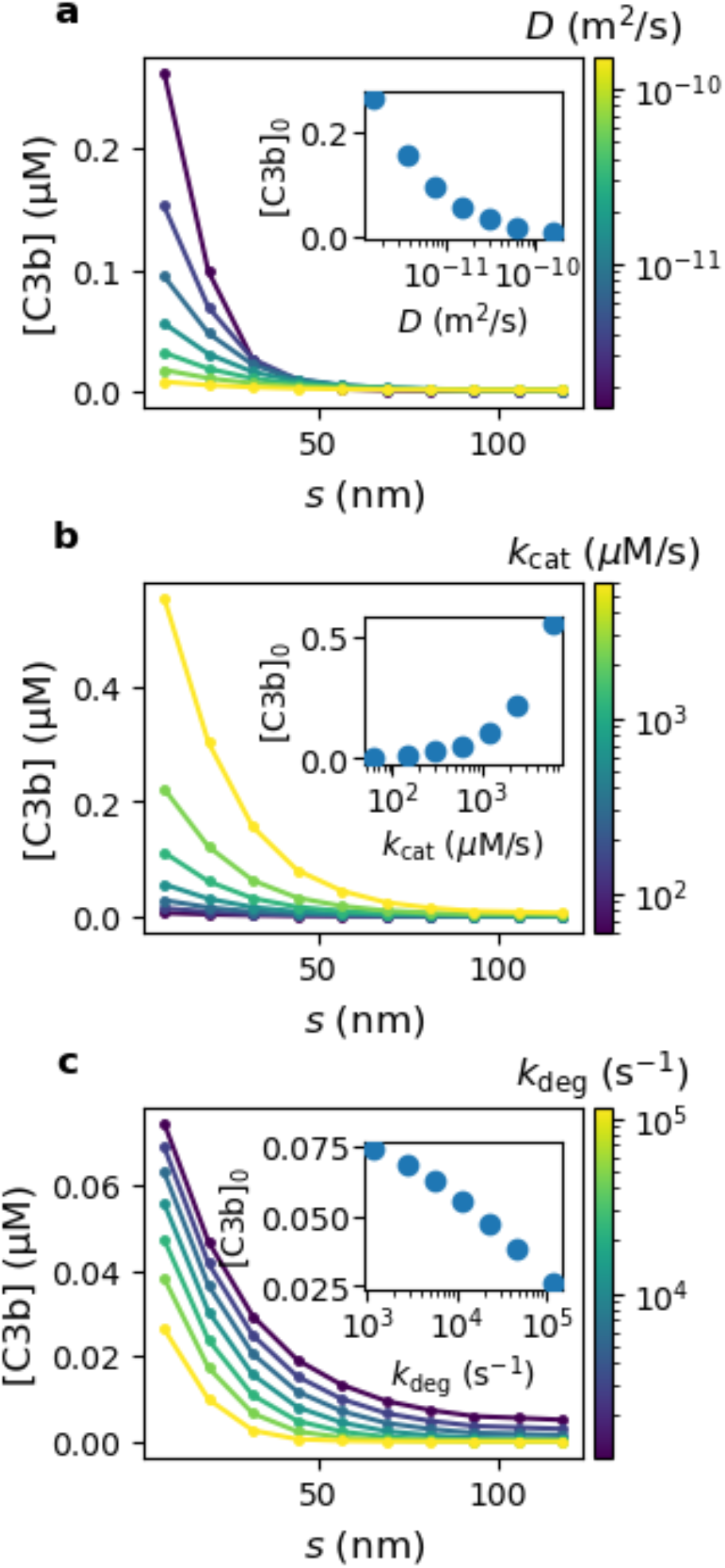
Spatial C3b concentration profile shifts with D (a), k_cat_ (b), and k_decay_ (c). Insets illustrate how the initial concentration value changes with the parameter value.

## Supplemental Tables

**Table S1:** Parameter values used in ABM simulations. *τ*_ABM_represents the simulation time unit. To map *τ*_ABM_ to seconds, we can solve using the experimentally observed half-life of C3b: 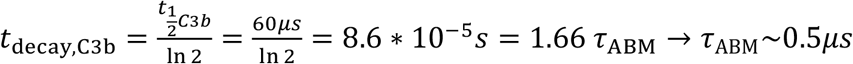

| ABM |  |  |
| --- | --- | --- |
| Parameter | Value | Source |
| Surface diffusion speed | $52.5 \frac{\text{nm}}{\tau_{ABM}}$ | Previous work<sup>1</sup> |
| Catalysis rate | $0.8 \frac{1}{\tau_{ABM}}$ | Previous work<sup>1</sup> |
| Decay time of bound C3b | $5.0 \tau_{ABM}$ | Previous work<sup>1</sup> |
| Decay time of unbound C3b | $1.66 \tau_{ABM}$ | Previous work<sup>1</sup> |

**Table S2:**
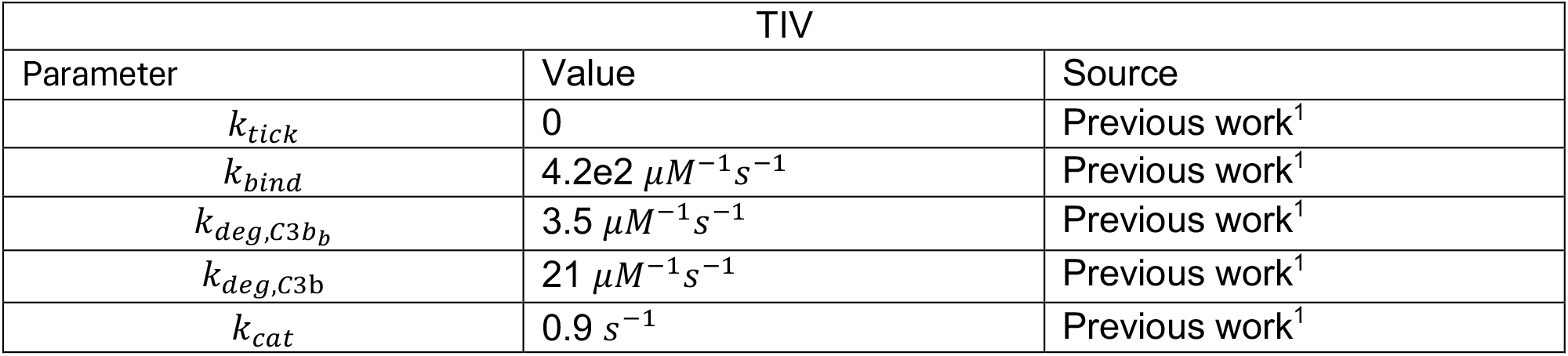
Parameter values used in TIV model.

| TIV |  |  |
| --- | --- | --- |
| Parameter | Value | Source |
| $k_{tick}$ | 0 | Previous work<sup>1</sup> |
| $k_{bind}$ | $4.2e2 \mu M^{-1} s^{-1}$ | Previous work<sup>1</sup> |
| $k_{deg,C3b_b}$ | $3.5 \mu M^{-1} s^{-1}$ | Previous work<sup>1</sup> |
| $k_{deg,C3b}$ | $21 \mu M^{-1} s^{-1}$ | Previous work<sup>1</sup> |
| $k_{cat}$ | $0.9 s^{-1}$ | Previous work<sup>1</sup> |

**Table S3:** Parameter values used in PDE model.

| PDE |  |  |
| --- | --- | --- |
| Parameter | Value | Source |
| $D_{C3b}$ | $2720 s^{-1}$ | C3b diffusion coefficient from literature $(1.53 * 10^{-11} \frac{m^2}{s})^2$, with length scale non-dimensionalized by nanoparticle radius (75 nm) |
| $k_{decay}$ | $1.6e4 s^{-1}$ | Computed using $t_{\frac{1}{2}\text{C3b}}$ from literature<sup>2</sup>. |
| $k_{cat,PDE}$ | $600 \frac{\mu M}{s}$ | Previous work<sup>1</sup>, renormalized for surface catalysis |
| $\sigma$ | 0.1 | Size of C3b protein (15 nm)<sup>3,4</sup>, non-dimensionalized by nanoparticle radius (75 nm). |
| $x_0$ | (0.0, 0.0, 1.2) | Size of C3b protein (15 nm)<sup>3,4</sup>, non-dimensionalized by nanoparticle radius (75 nm). |

## Supplemental Text

### S1 Empirical fit for critical exponents

We first fit a cubic spline interpolation to the collected data points using the CubicSpline function from SciPy so that we can fit our power law curve close to the critical point^5^. We also identify the critical point as *θ*_*c*_ at max(*M*_*τ*_) according to critical transition theory. We define the reduced distance as 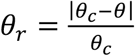. We now fit the following functions using points 0.01 < *θ*_*r*_ < 0.1 with *θ* on the activated side of the transition

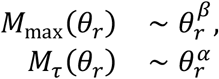

using SciPy’s curve_fit function^5^.

### S2 Mean-field derivation of the critical exponents

#### S2.1 Defining threshold conditions for mean-field system

We first seek to define the control parameter at the threshold position. To do so, we write the Jacobian for C3b and C3bb for the activation-free state ([Sites], [C3b], [C3b_b_]) = ([Sites]_0_, 0,0), where [Sites]_0_ is the initial concentration of available sites:

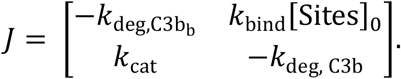

This gives the characteristic equation

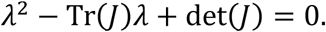

At a critical point, one of the roots is *λ* = 0. Thus, det(*J*) = 0, and we can solve

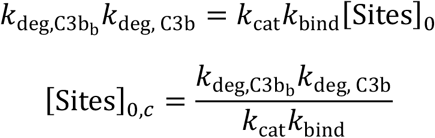

Here, [Sites]_0_ is the control parameter, related to site spacing as [Sites]_0_ ∝ *d*^−2^, and [Sites]_0,*c*_is the critical point. We define the reduced control parameter

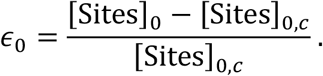

These definitions will be used for computing the critical exponents in the following subsections.

#### S2.2 Computing lifetime exponent near threshold

The roots of the characteristic equation of *J* at exactly the critical threshold *λ* = 0 and 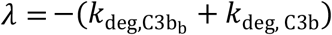. The slow eigenvalue, *λ* = 0, represents the divergence of the activity lifetime at the critical threshold. To compute the timescale of this slow mode when approaching the critical threshold, we define *λ*_+_as a small perturbation to *λ* above 0.

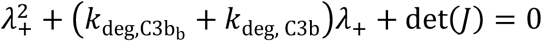

Since 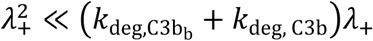, we keep only first-order terms. We write 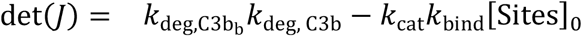 in terms of *ϵ* using the relations above, giving 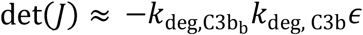 near the threshold. This gives

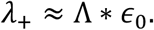

where Λ is a constant defined by

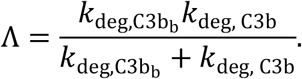

The characteristic activation timescale is the inverse 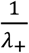, giving 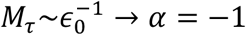.

This agrees with our empirical fit and the dynamic percolation timescale described for the general epidemic process in literature^6^.

#### S2.3 Computing peak-activity exponent near threshold

We next aim to define the peak activity as a function of *ϵ*.

Recall

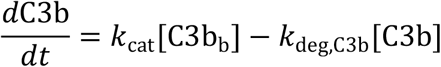

and

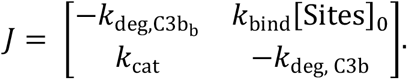

From the eigenvector associated with *λ* = 0, we find 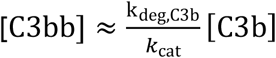 near the threshold.

Thus,

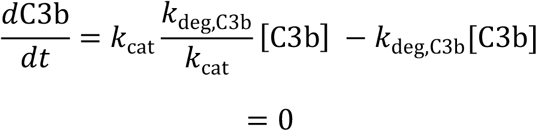

at the threshold. Perturbations slightly above the threshold give

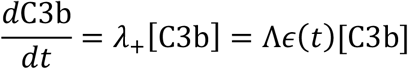

where

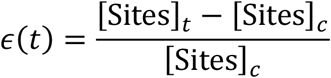

and

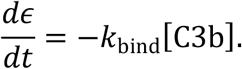

Writing how C3b changes as a function of *ϵ*:

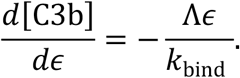

Integrating gives

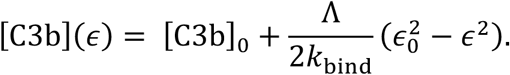

The peak activity occurs where 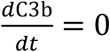, which is also the point where available [Sites]_*t*_ crosses the threshold, and thus *ϵ* = 0. For a small [C3b]_0_, as is the case in our system with a small tickover seed, 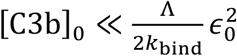. Thus,

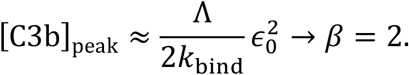

This agrees with our empirical fit. As mentioned in the main text, there has not yet been much analysis of this quantity in previous literature and thus there does not appear to be a directly published mean-field counterpart.

Thus, both empirical locus exponents follow from closed-form analysis of the reaction network itself.

## Notes

### Competing Interest Statement

The authors have declared no competing interest.

## References

1 J.M. Anderson, A. Rodriguez, and D.T. Chang, “Foreign body reaction to biomaterials,” Seminars in Immunology 20(2), 86–100 (2008).

2 Q. Chen, D. Zhang, J. Gu, H. Zhang, X. Wu, C. Cao, X. Zhang, and R. Liu, “The impact of antifouling layers in fabricating bioactive surfaces,” Acta Biomaterialia 126, 45–62 (2021).

3 R. Cai, and C. Chen, “The Crown and the Scepter: Roles of the Protein Corona in Nanomedicine,” Advanced Materials 31(45), 1805740 (2019).

4 S.M. Moghimi, A.J. Andersen, D. Ahmadvand, P.P. Wibroe, T.L. Andresen, and A.C. Hunter, “Material properties in complement activation,” Advanced Drug Delivery Reviews 63(12), 1000–1007 (2011).

5 S.M. Moghimi, D. Simberg, E. Papini, and Z.S. Farhangrazi, “Complement activation by drug carriers and particulate pharmaceuticals: Principles, challenges and opportunities,” Advanced Drug Delivery Reviews 157, 83–95 (2020).

6 S.M. Moghimi, H.B. Haroon, A. Yaghmur, D. Simberg, and P.N. Trohopoulos, “Nanometer- and angstrom-scale characteristics that modulate complement responses to nanoparticles,” Journal of Controlled Release 351, 432–443 (2022).

7 J. Szebeni, D. Simberg, Á. González-Fernández, Y. Barenholz, and M.A. Dobrovolskaia, “Roadmap and strategy for overcoming infusion reactions to nanomedicines,” Nat Nanotechnol 13(12), 1100–1108 (2018).

8 Kenneth Murphy and Casey Weaver, Janeway’s Immunobiology, 9th edition (W.W. Norton & Company, New York, 2017).

9 A. Gorter, S. Meri, A. Gorter, and S. Meri, “Immune evasion of tumor cells using membrane-bound complement regulatory proteins,” Immunology Today 20(12), 576–582 (1999).

10 A.J. Tenner, and T.J. Petrisko, “Knowing the enemy: strategic targeting of complement to treat Alzheimer disease,” Nat Rev Neurol 21(5), 250–264 (2025).

11 D. Ricklin, “Complement-targeted therapeutics: Are we there yet, or just getting started?,” Eur J Immunol 54(12), 2350816 (2024).

12 E.S. Reis, D.C. Mastellos, D. Ricklin, A. Mantovani, and J.D. Lambris, “Complement in cancer: untangling an intricate relationship,” Nat Rev Immunol 18(1), 5–18 (2018).

13 K. Peters, “Physiology and pathology of the C3 amplification cycle: A retrospective,” Immunological Reviews 313(1), 217–224 (2023).

14 P.J. Lachmann, “Looking back on the alternative complement pathway,” Immunobiology 223(8), 519–523 (2018).

15 P.J. Lachmann, “The Amplification Loop of the Complement Pathways,” in Advances in Immunology, (Academic Press, 2009), pp. 115–149.

16 M.K. Pangburn, “Initiation of the alternative pathway of complement and the history of ‘tickover,’” Immunological Reviews 313(1), 64–70 (2023).

17 Z. Ban, P. Yuan, F. Yu, T. Peng, Q. Zhou, and X. Hu, “Machine learning predicts the functional composition of the protein corona and the cellular recognition of nanoparticles,” Proceedings of the National Academy of Sciences 117(19), 10492–10499 (2020).

18 Z. Wang, S. Kulkarni, J. Nong, M. Zamora, A. Ebrahimimojarad, E. Hood, T. Shuvaeva, M. Zaleski, D. Gullipalli, E. Wolfe, C. Espy, E. Arguiri, J. Wu, Y. Wang, O.A. Marcos-Contreras, W. Song, V.R. Muzykantov, J. Fu, R. Radhakrishnan, J.W. Myerson, and J.S. Brenner, “A percolation phase transition controls complement protein coating of surfaces,” Cell 188(15), 4058–4073.e25 (2025).

19 P. Grassberger, “On the critical behavior of the general epidemic process and dynamical percolation,” Mathematical Biosciences 63(2), 157–172 (1983).

20 D. Stauffer, and A. Aharony, Introduction To Percolation Theory: Second Edition, 2nd ed. (Taylor & Francis, London, 2018).

21 X.-L. Peng, S.-Y. Chang, S. Chen, and G.-Q. Sun, “Percolation thresholds and epidemic spreading on small-world networks: Exact results and critical dynamics,” Mathematics and Computers in Simulation 245, 192–211 (2026).

22 M. Kivelä, J. Cambe, J. Saramäki, and M. Karsai, “Mapping temporal-network percolation to weighted, static event graphs,” Sci Rep 8(1), 12357 (2018).

23 J. Almeira, D.A. Martin, D.R. Chialvo, and S.A. Cannas, “Susceptibility for extremely low external fluctuations and critical behavior of Greenberg-Hastings neuronal model,” Phys. Rev. E 113(2), 024311 (2026).

24 C. Autorino, and N.I. Petridou, “Critical phenomena in embryonic organization,” Current Opinion in Systems Biology 31, 100433 (2022).

25 J.W. Larkin, X. Zhai, K. Kikuchi, S.E. Redford, A. Prindle, J. Liu, S. Greenfield, A.M. Walczak, J. Garcia-Ojalvo, A. Mugler, and G.M. Süel, “Signal Percolation within a Bacterial Community,” Cels 7(2), 137–145.e3 (2018).

26 M.A. Aon, S. Cortassa, and B. O’Rourke, “Percolation and criticality in a mitochondrial network,” Proc Natl Acad Sci U S A 101(13), 4447–4452 (2004).

27 N. Zewde, R.D. Gorham, A. Dorado, and D. Morikis, “Quantitative Modeling of the Alternative Pathway of the Complement System,” PLoS ONE 11(3), e0152337 (2016).

28 J.M. Yeomans, Statistical Mechanics of Phase Transitions (Oxford University Press, Oxford, New York, 1992).

29 S.T. Brown, P. Buitrago, E. Hanna, S. Sanielevici, R. Scibek, and N.A. Nystrom, “Bridges-2: A Platform for Rapidly-Evolving and Data Intensive Research,” in Practice and Experience in Advanced Research Computing 2021: Evolution Across All Dimensions, (Association for Computing Machinery, New York, NY, USA, 2021), pp. 1–4.

30 P. Baccam, C. Beauchemin, C.A. Macken, F.G. Hayden, and A.S. Perelson, “Kinetics of Influenza A Virus Infection in Humans,” Journal of Virology 80(15), 7590–7599 (2006).

31 C. Rackauckas, and Q. Nie, “DifferentialEquations.jl – A Performant and Feature-Rich Ecosystem for Solving Differential Equations in Julia,” JORS 5(1), 15 (2017).

32 S. Badia, and F. Verdugo, “Gridap: An extensible Finite Element toolbox in Julia,” JOSS 5(52), 2520 (2020).

33 F. Verdugo, and S. Badia, “The software design of Gridap: A Finite Element package based on the Julia JIT compiler,” Computer Physics Communications 276, 108341 (2022).

34 C.A. Browne, D.B. Amchin, J. Schneider, and S.S. Datta, “Infection Percolation: A Dynamic Network Model of Disease Spreading,” Front. Phys. 9, (2021).

35 T. Mora, and W. Bialek, “Are Biological Systems Poised at Criticality?,” J Stat Phys 144(2), 268–302 (2011).

36 C. Trefois, P.M. Antony, J. Goncalves, A. Skupin, and R. Balling, “Critical transitions in chronic disease: transferring concepts from ecology to systems medicine,” Current Opinion in Biotechnology 34, 48–55 (2015).

37 N. Rappaport, B. Nogal, K. Perrott, V. Domina, L. Hood, and N.D. Price, “Early Detection of Wellness-to-Disease Transitions in the AI Era: Implications for Pharmacology and Toxicology,” Annual Review of Pharmacology and Toxicology 66(Volume 66, 2026), 41–64 (2026).

38 A. Presbitero, R. Quax, V.V. Krzhizhanovskaya, and P.M.A. Sloot, “Detecting Critical Transitions in the Human Innate Immune System Post-cardiac Surgery,” Computational Science – ICCS 2020 12137, 371–384 (2020).

39 Z. Wang, E.D. Hood, J. Nong, J. Ding, O.A. Marcos-Contreras, P.M. Glassman, K.M. Rubey, M. Zaleski, C.L. Espy, D. Gullipali, T. Miwa, V.R. Muzykantov, W.-C. Song, J.W. Myerson, and J.S. Brenner, “Combating Complement’s Deleterious Effects on Nanomedicine by Conjugating Complement Regulatory Proteins to Nanoparticles,” Advanced Materials 34(8), 2107070 (2022).

40 Y. Li, S. Jacques, H. Gaikwad, G. Wang, N.K. Banda, V.M. Holers, R.I. Scheinman, S. Tomlinson, S.M. Moghimi, and D. Simberg, “Inhibition of acute complement responses towards bolus-injected nanoparticles using targeted short-circulating regulatory proteins,” Nat. Nanotechnol. 19(2), 246–254 (2024).

41 Y. Li, S. Jacques, H. Gaikwad, M. Nebbia, N.K. Banda, V.M. Holers, S.A. Tomlinson, R.I. Scheinman, A. Monte, L. Saba, E. Lasda, J. Hasselberth, N. Busquet, W.M. Zelek, S.M. Moghimi, and D. Simberg, “Enhanced immunocompatibility and hemocompatibility of nanomedicines across multiple species using complement pathway inhibitors,” Science Advances 11(28), eadw1731 (2025).

42 J.H. Park, J.A. Jackman, A.R. Ferhan, J.N. Belling, N. Mokrzecka, P.S. Weiss, and N.-J. Cho, “Cloaking Silica Nanoparticles with Functional Protein Coatings for Reduced Complement Activation and Cellular Uptake,” ACS Nano 14(9), 11950–11961 (2020).

## Supplementary References

1 Z. Wang, S. Kulkarni, J. Nong, M. Zamora, A. Ebrahimimojarad, E. Hood, T. Shuvaeva, M. Zaleski, D. Gullipalli, E. Wolfe, C. Espy, E. Arguiri, J. Wu, Y. Wang, O.A. Marcos-Contreras, W. Song, V.R. Muzykantov, J. Fu, R. Radhakrishnan, J.W. Myerson, and J.S. Brenner, “A percolation phase transition controls complement protein coating of surfaces,” Cell 188(15), 4058–4073.e25 (2025).

2 N. Zewde, R.D. Gorham, A. Dorado, and D. Morikis, “Quantitative Modeling of the Alternative Pathway of the Complement System,” PLoS ONE 11(3), e0152337 (2016).

3 B.J.C. Janssen, A. Christodoulidou, A. McCarthy, J.D. Lambris, and P. Gros, “Human Complement Component C3b: 2i07,” (2006).

4 B.J.C. Janssen, A. Christodoulidou, A. McCarthy, J.D. Lambris, and P. Gros, “Structure of C3b reveals conformational changes that underlie complement activity,” Nature 444(7116), 213–216 (2006).

5 P. Virtanen, R. Gommers, T.E. Oliphant, M. Haberland, T. Reddy, D. Cournapeau, E. Burovski, P. Peterson, W. Weckesser, J. Bright, S.J. Van Der Walt, M. Brett, J. Wilson, K.J. Millman, N. Mayorov, A.R.J. Nelson, E. Jones, R. Kern, E. Larson, C.J. Carey, İ. Polat, Y. Feng, E.W. Moore, J. VanderPlas, D. Laxalde, J. Perktold, R. Cimrman, I. Henriksen, E.A. Quintero, C.R. Harris, A.M. Archibald, A.H. Ribeiro, F. Pedregosa, P. Van Mulbregt, SciPy 1.0 Contributors, A. Vijaykumar, A.P. Bardelli, A. Rothberg, A. Hilboll, A. Kloeckner, A. Scopatz, A. Lee, A. Rokem, C.N. Woods, C. Fulton, C. Masson, C. Häggström, C. Fitzgerald, D.A. Nicholson, D.R. Hagen, D.V. Pasechnik, E. Olivetti, E. Martin, E. Wieser, F. Silva, F. Lenders, F. Wilhelm, G. Young, G.A. Price, G.-L. Ingold, G.E. Allen, G.R. Lee, H. Audren, I. Probst, J.P. Dietrich, J. Silterra, J.T. Webber, J. Slavič, J. Nothman, J. Buchner, J. Kulick, J.L. Schönberger, J.V. De Miranda Cardoso, J. Reimer, J. Harrington, J.L.C. Rodríguez, J. Nunez-Iglesias, J. Kuczynski, K. Tritz, M. Thoma, M. Newville, M. Kümmerer, M. Bolingbroke, M. Tartre, M. Pak, N.J. Smith, N. Nowaczyk, N. Shebanov, O. Pavlyk, P.A. Brodtkorb, P. Lee, R.T. McGibbon, R. Feldbauer, S. Lewis, S. Tygier, S. Sievert, S. Vigna, S. Peterson, S. More, T. Pudlik, T. Oshima, T.J. Pingel, T.P. Robitaille, T. Spura, T.R. Jones, T. Cera, T. Leslie, T. Zito, T. Krauss, U. Upadhyay, Y.O. Halchenko, and Y. Vázquez-Baeza, “SciPy 1.0: fundamental algorithms for scientific computing in Python,” Nat Methods 17(3), 261–272 (2020).

6 P. Grassberger, “On the critical behavior of the general epidemic process and dynamical percolation,” Mathematical Biosciences 63(2), 157–172 (1983).

